# Systematic protein complex profiling and differential analysis from co-fractionation mass spectrometry data

**DOI:** 10.1101/2020.05.06.080465

**Authors:** Andrea Fossati, Chen Li, Peter Sykacek, Moritz Heusel, Fabian Frommelt, Federico Uliana, Mahmoud Hallal, Isabell Bludau, Capraz Tümay Klemens, Peng Xue, Anthony W. Purcell, Matthias Gstaiger, Ruedi Aebersold

## Abstract

Protein complexes, macro-molecular assemblies of two or more proteins, play vital roles in numerous cellular activities and collectively determine the cellular state. Despite the availability of a range of methods for analysing protein complexes, systematic analysis of complexes under multiple conditions has remained challenging. Approaches based on biochemical fractionation of intact, native complexes and correlation of protein profiles have shown promise, for instance in the combination of size exclusion chromatography (SEC) with accurate protein quantification by SWATH/DIA-MS. However, most approaches for interpreting co-fractionation datasets to yield complex composition, abundance and rearrangements between samples depend heavily on prior evidence. We introduce PCprophet, a computational framework to identify novel protein complexes from SEC-SWATH-MS data and to characterize their changes across different experimental conditions. We demonstrate accurate prediction of protein complexes (AUC >0.99 and accuracy around 97%) via five-fold cross-validation on SEC-SWATH-MS data, show improved performance over state-of-the-art approaches on multiple annotated co-fractionation datasets, and describe a Bayesian approach to analyse altered protein-protein interactions across conditions. PCprophet is a generic computational tool consisting of modules for data pre-processing, hypothesis generation, machine-learning prediction, post-prediction processing, and differential analysis. It can be applied to any co-fractionation MS dataset, independent of separation or quantitative LC-MS workflow employed, and to support the detection and quantitative tracking of novel protein complexes and their physiological dynamics.

## Main

The analysis of proteins has progressed from studying specific proteins to the comparative analysis of multiple proteomes, allowing for the detection of changes in the proteome landscape as a function of the cellular state and the identification of connections between proteins based on their behaviours across multiple samples. However, proteins largely function as complexes which are involved in performing and regulating a majority of biological functions^1–4^. Protein complexes are a part of extended functional groups, such as pathways or protein interaction networks. Despite the availability of a range of methods for the analysis of specific protein complexes, systematic analysis of the ensemble of protein complexes in a sample has remained challenging^5^. While affinity-purification mass spectrometry (AP-MS) provides valuable biological information on protein complexes, it lacks scalability and requires either genetic manipulation of cells for introduction of a tag or the use of antibody-based reagents. On the other hand, biochemical fractionation mass spectrometry allows for simultaneous quantification of thousands of proteins and is emerging as a powerful technique for system-wide investigation of protein complexes. Analytical techniques such as size exclusion chromatography (SEC) and ion exchange chromatography (IEX) have been successfully applied in a variety of complex biological questions such as apoptosis-dependent complex rewiring^6^, characterization of novel complexes in *Trypanosoma Brucei*^7^ and *C. Elegans*^8^, identification of isoform-specific complexes^9^ and differential analysis of cell cycle states^17^, i.e. the interphase and mitosis.

A key challenge in fractionation-based approaches is the is the confident assignment of protein subunits to protein complexes based on their co-fractionation patterns and other relevant biological information. A number of computational frameworks have been proposed for this purpose^8, 10–12^. Among these methods, CCprofiler identifies protein complexes from co-fractionation proteomic data based on prior information from reference complex/interactome databases such as CORUM^13^, STRING^14^ and BioPlex^15, 16^. CCprofiler was not designed to predict novel protein complexes but to determine a confidently detectable set of complexes including statistical estimation and control of the false discovery rate^10^. PrInCE and EPIC leverage machine-learning techniques to predict novel protein complexes but are limited conceptually to the inference of protein-protein interactions (PPI) from co-fractionation proteomic data. Finally, dendrogram clustering has been described for novel complex identification^11^. In this case as well, control of false positives and false negatives is challenging since an arbitrary threshold must be applied to cut the dendrogram^11^.

In this study, we describe PCprophet, an open-access software for protein complex prediction directly from co-fractionation-MS data using machine-learning techniques and differential analysis of complex abundance and assembly state across conditions. PCprophet combines the benefits from previous approaches such as error rate control using database-derived complexes present in CCprofiler with the discovery of novel complexes inherent in other approaches. PCprophet offers the following features: (i) PCprophet accepts input from a variety of co-fractionation mass-spectrometry (coFrac-MS) techniques, including but not limited to size-exclusion chromatography (SEC-MS), strong cationic exchange (SCX), and blue native page (BNP); (ii) PCprophet can be used with inputs derived from widely employed mass spectrometry acquisition schemes such as data dependent acquisition (DDA), data independent acquisition (DIA) and different quantitation strategies such isobaric labelling (SILAC, TMT) or label-free; (iii) PCprophet was trained using co-eluting protein complex data, rather than co-eluting PPIs, and can therefore directly predict novel protein complexes (i.e. complex-centric prediction); (iv) PCprophet performs post-prediction processing via a statistical error model based on Gene Ontology scores and other criteria to improve the reliability of the predicted protein complexes and reduce false positives; (v) PCprophet performs differential analysis of predicted protein complexes across conditions using our newly proposed Bayesian inference-based method. We applied PCprophet to predict and analyse protein complexes in different cell cycle phases using our recently published SEC-SWATH-MS dataset in the HeLa cell line^17^. Our results demonstrate that PCprophet predicts novel protein complexes and recapitulates known changes in protein complexes across the cell cycle.

## Results

### PCprophet accurately identifies novel protein complexes from co-fractionation MS data

PCprophet enables accurate prediction of protein complexes directly from raw input (i.e. protein matrices consisting of protein intensity vs. fraction number) of SEC-SWATH-MS and other co-fractionation data. The framework of PCprophet (**Fig. 1**) includes six major modules: data pre-processing, database query and *de novo* complex (i.e. hypothesis) generation, feature calculation and prediction, error estimation and post-prediction processing, complex-centric differential analysis, and report generation and data visualisation. During the data pre-processing step, Gaussian filtering, missing value imputation, linear interpolation and data resizing are performed to ensure data quality (See ‘**Methods**’ for more details). During the hypothesis generation step, a list of candidate protein complexes for each condition based on the raw input protein matrices is provided separately via peak-picking and distance-based clustering, for the machine-learning model to predict. During feature calculation and prediction, each protein complex delivered by the hypothesis generation procedure is represented using a numeric vector, including average intensity difference of proteins within each fraction, local correlation of proteins at each window, shift of apex fraction of each protein and average full width of a peak at half maximum. Meanwhile, the provided database (either PPI or complexes) is mapped in the same feature space for later being used for FDR control. Then the Random Forest models predict potential protein complexes with detailed predicted probabilities. During error estimation and post-prediction processing, PCprophet filters the predictions based on Gene Ontology (GO) terms assigned to components of predicted complexes. By calculating the pairwise GO term semantic similarities of proteins assigned to a complex and comparing them to similarity scores in reference databases of known protein complexes, PCprophet filters predicted complexes by a local false discovery rate (FDR) based on GO term semantic similarity. In addition, PCprophet performs complex combination and collapsing, since hypothesized complexes might be a subset of a bigger complex or a mix of multiple complexes. During the complex-centric differential analysis, PCprophet analyses the differences in prediction results between conditions, from protein level to complex level, using a Bayesian inference method. As the final output, tabular and visual reports of the predicted protein complexes and their changes across different conditions are generated by PCprophet. In summary, PCprophet provides a ‘one-stop’ computational framework for the confident detection of protein complexes including their dynamic changes across different biological states from a wide range of coFrac-MS data.

**Fig. 1.**
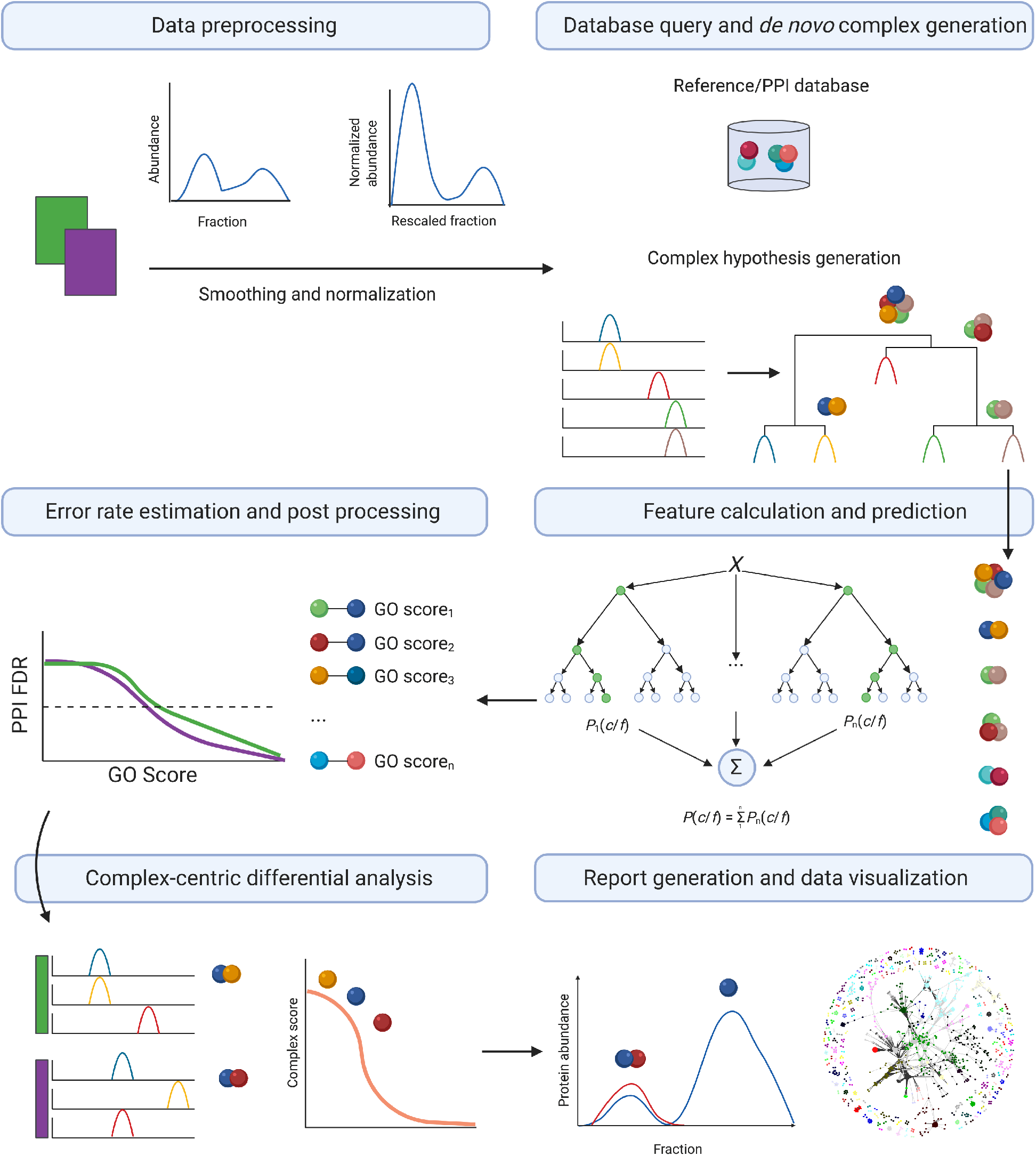
The framework of PCprophet. It consists of the six major modules including (i) data pre-processing, (ii) database query and *de novo* complex (i.e. hypothesis) generation, (iii) feature calculation and prediction, (iv) error estimation and post-prediction processing, (v) complex-centric differential analysis, and (vi) report generation and data visualisation.

### Benchmarking PCprophet complex prediction against state-of-the-art methods

Concluding from the five-fold cross-validation (refer to ‘**Supplementary Results**’ for more details), Random Forest (RF) has been chosen as the core classification algorithm of PCprophet. We then assessed the performance of complex predictions using the optimized PCprophet framework against two different, state-of-the-art computational approaches for the detection of protein complexes from co-fractionation data, namely CCprofiler^10^ and EPIC^8^. Similar to PrInCE^12^, EPIC supports the prediction of binary protein-protein interactions and network inference of underlying complexes, maintaining the potential to discover previously unknown complexes. Out of these two interaction-centric approaches, we selected EPIC for our performance comparison as it has been shown to outperform previous tools such as PrInCE. We benchmarked these tools based on a recently published dataset, where HeLa CCL2 cells were synchronized in distinct cell-cycle stages (i.e. interphase and mitosis) prior to analysis by SEC and DIA/SWATH-MS^17^. To avoid biases in assessing performance due to the different inputs required by these tools (CCprofiler mainly takes the peptide-level quantitative values as input, whereas PCprophet and EPIC take as input the protein-level quantitative values), we performed sibling peptide correlation using CCprofiler and exported the resulting protein matrices, thereby providing the same input for all benchmarked tools (refer to ‘**Methods’** for more details). To minimize comparison bias due to parameter optimization, we ran CCprofiler with the parameters used in its original publication^10^. EPIC, on the other hand, offers the possibility of choosing between an SVM classifier or an RF classifier for PPI prediction. We used default parameters with both classifiers, generating two sets of predictions (EPIC_SVM and EPIC_RF). We generated protein complex hypotheses for CCprofiler using the CORUM core complexes dataset, and also trained EPIC using CORUM. PCprophet requires a protein complex or PPI database as input to perform FDR control and CORUM was used for this purpose as well.

We initially evaluated the performance of each method using the numbers of known CORUM complexes recovered across all replicates and conditions. Both the absolute number of identified complexes as well as the overall recall are vastly different for each tool. The complex-centric tools (i.e. PCprophet and CCprofiler) identified 900 and 798 known complexes respectively; while EPIC_RF recovered only 71 known complexes and EPIC_SVM recovered none (**Fig. 2a**). The overlap in known complexes between PCprophet and CCprofiler was 69.7% (i.e. 556 out of 798), while 49.2% (i.e. 35 out of 71) overlap was achieved between PCprophet and EPIC_RF (**Fig. 2b**). The identified complexes correspond to a recall rate of 37% for PCprophet, 33% for CCprofiler and 3% for EPIC_RF. PCprophet recalls a much higher fraction of CORUM complexes compared to those recalled by EPIC analysis also in a DDA-based dataset with a isotope dilution strategy for quantification^18^ (DDA-SILAC, **Supplementary Fig. S1**). We then compared the average number of subunits per complex to evaluate the similarity of known complexes from CORUM to the predicted ones from EPIC and PCprophet (**Fig. 2c**). The distribution of subunits per complex predicted by PCprophet and CCprofiler is closer to that of CORUM complexes (average 4.1 subunits), with an average subunit size of 3.5 for PCprophet and 6.9 for CCprofiler. The average subunit size per complex predicted by EPIC, however, was 19.9 (p<10E-14) for the RF classifier and 71.7 (p<10E-14) for the SVM classifier, respectively. The results from EPIC thus suggest a larger size of cellular assemblies compared to the sizes of manually curated complexes in the CORUM database, with more similar sizes reported by both CCprofiler and PCprophet.

**Fig. 2.**
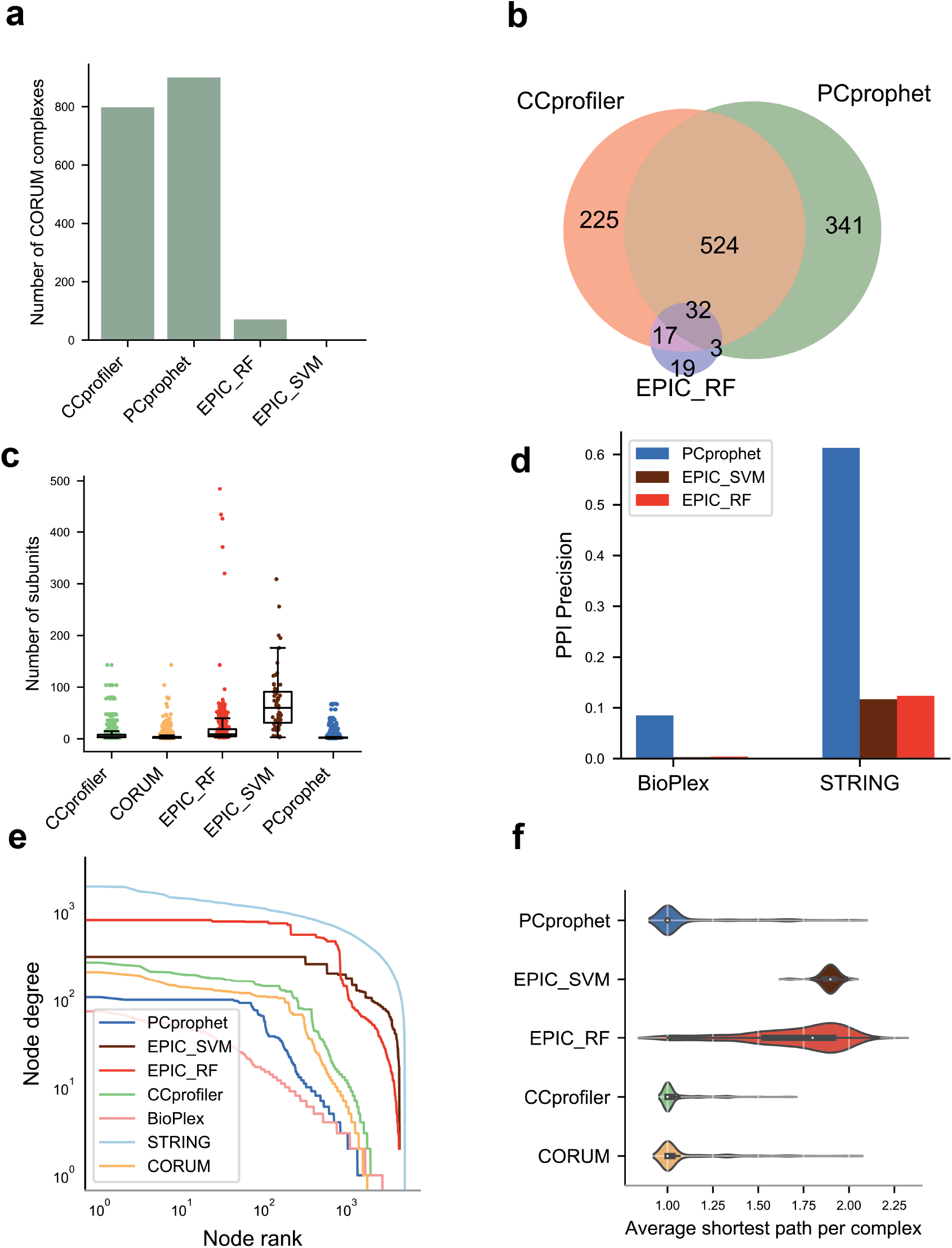
Benchmarking PCprophet against existing tools for protein complex profiling and prediction. **a**, The numbers of CORUM complexes recovered and the numbers of overlapping complexes by the assessed tools. **b**, Absolute number of CORUM complexes recovered by each tool. **c**, Number of subunits per complex predicted and identified by different tools. Boxplot shows the medians and the ticks represent standard deviation. **d**, The precision values (refer to the ‘Methods’ section for more details) of PPI prediction for *de novo* protein complex prediction tools. **e**, Log-log plot showing the degree distribution of the network generated by each tool *versus* ground-truth databases (STRING, BioPlex and CORUM). **f**, Distribution of shortest path per complex across all subunits, as reported by the indicated tools. The medians are highlighted in white dots.

We then evaluated the performance of PCprophet and EPIC in recalling protein-protein interactions (PPIs). In this comparison, we did not consider CCprofiler as it cannot derive novel complexes without prior information, which limits its applicability for discovery of novel protein-protein interactions. We generated a PPI network from complexes predicted by PCprophet, EPIC_RF, and EPIC_SVM, and compared them to ground truth networks from CORUM complexes and from PPI databases such as STRING and BioPlex. This comparison allows to calculate the percentage of reported PPIs for each tool, in the form of PPI precision. PCprophet achieved a PPI precision of 0.65 when compared with STRING and 0.095 in comparison to BioPlex database (**Fig. 2d**). The precision of EPIC_RF was 0.12 with STRING and 0.004 when compared with PPIs in BioPlex. EPIC_SVM prediction corresponded to a precision of 0.11 and 0.002 with STRING and BioPlex respectively. We calculated for each network the degree distribution and the frequency of nodes with a particular degree (**Fig. 2e**). To evaluate the similarity between ground-truth networks and prediction, the Area Under the Curve (AUC) values were calculated for all the tools (**Supplementary Table S1**) and databases. Regardless of the classifier used, EPIC-derived PPI networks tend to have higher degree (**Fig. 2e**) compared to those from complexes in CORUM. This resulted in an AUC of 0.18 for EPIC_RF, 0.39 for EPIC_SVM, 0.13 for PCprophet and 0.11 for CORUM, respectively. In this context, an AUC value closer to the one of reported complexes (CORUM) means a closer resemblance in network topology to a ground truth network. Finally, we merged all PPI databases (STRING, BioPlex and BioGrid) to generate a combined network including all deposited interactions and assessed the average distance between every pair of proteins within a predicted or known complex. The average shortest path for EPIC_RF was 2.3 and >3 for EPIC_SVM while PCprophet-predicted complexes had an average path of 1.1 edges as shown in **Fig. 2f**, suggesting a greater recovery of closely connected proteins by PCprophet when compared to the average shortest path in CORUM (1 edge). We also observed a similar trend on an independent dataset from PrInCE^12^ (**Supplementary Fig. S1**). To summarize, PCprophet allows for robust identification of complexes, as shown based on high recall of known complexes and high quality of newly predicted complexes, demonstrated based on the high validation rates of the underlying PPIs by large-scale databases. PCprophet outperforms available tools in the recovery of known protein complexes and PPIs while additionally providing the opportunity to detect, investigate and track assemblies that remained inaccessible to computational approaches limited by their dependence on prior knowledge^10^ or the sensitivity of interaction-centric scoring^8, 12, 19^. In order to control spurious co-elution and false positive assignments, we integrated an error model based on interactor gene ontology similarity which effectively ensures highest quality of the reported results (**Supplementary Fig. S2**).

### Predicting PPIs and protein complexes across the mammalian cell cycle via PCprophet

We applied PCprophet to a second, newly published dataset^17^ in which HeLa cells were blocked at mitosis and interphase stages of the cell cycle. Proteins were then extracted under native conditions, SEC separated into 65 fractions and analysed using SWATH-MS. Based on these data, we generated a large PPI map based on all PCprophet predictions (**Fig. 3a**).

**Fig. 3.**
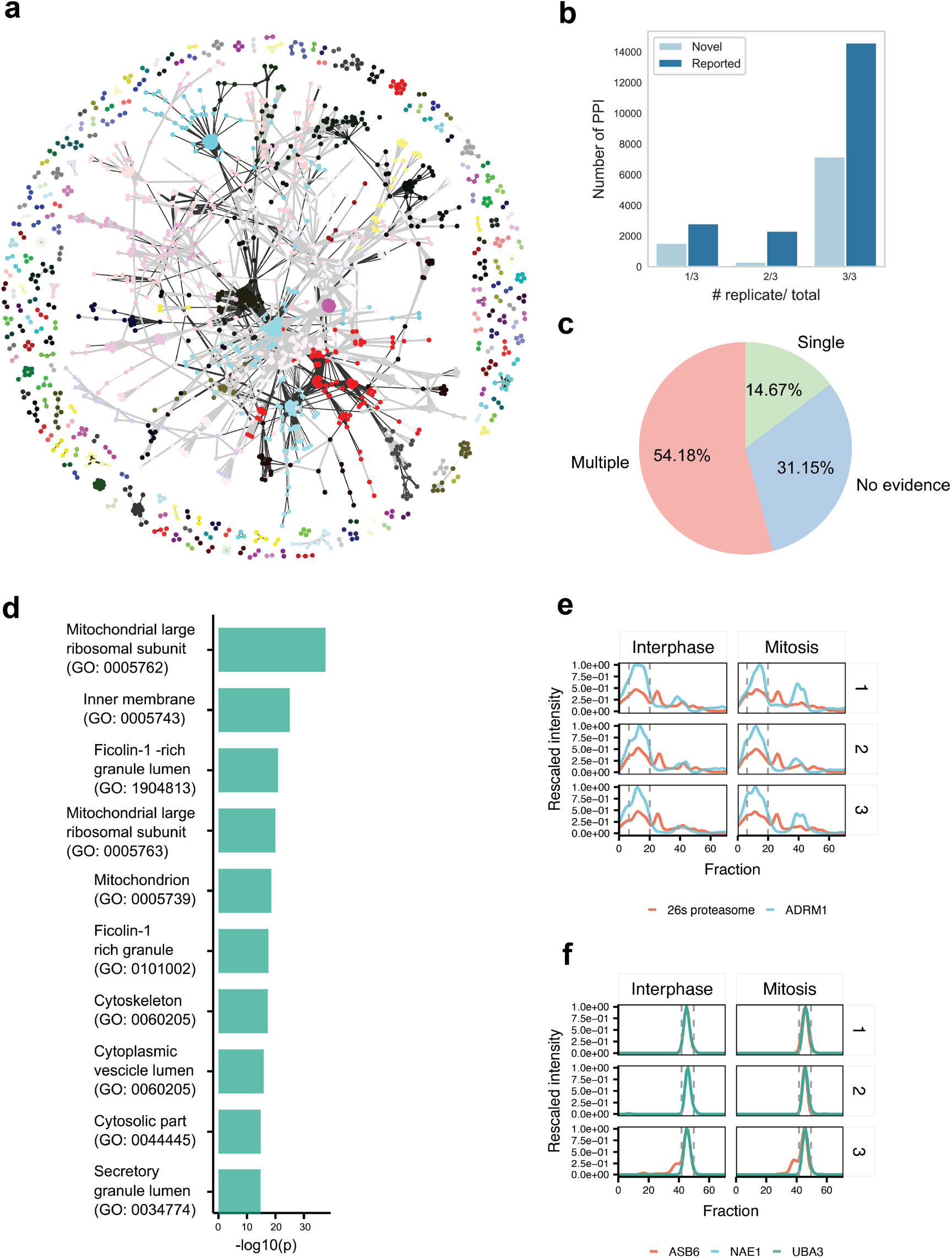
Evaluation of *de novo* prediction using PCprophet. **a**, The PPI map generated by PCprophet from HeLa cell proteomic data. Edge width represents the number of technical replicates for which a particular PPI was found. Black edge are novel PPIs and grey edges are reported PPIs. Protein communities are highlighted in different node colours. **b**, Number of novel and reported PPIs across all conditions within technical replicates (i.e. interphase and mitosis). **c**, Annotation of novel PPIs (i.e. not documented in the CORUM databases) in PPI databases (STRING, BioPlex, BioGrid). **d**, Enrichment analysis for GO Cellular Component for PPIs without prior evidence in any database. **e**, 26S proteasome and ADRM1 coelution from interphase and mitotic cells. **f**, Co-elution of ASB6 with the NAE1 and UBA3 complex.

PCprophet predicted 858 protein complexes not recorded in the CORUM derived network, which contain 11527 unique PPIs consistently present across all biological replicates for one condition (**Fig. 3b**), suggesting good reproducibility across different fractionation experiments. Of these predicted PPIs, 54.16% are consistently supported by evidence across several PPI databases (STRING, BioPlex and BioGrid^20^), 14.67% have PPI evidence from a single database while 31.14 % of the PPIs are completely novel (**Fig. 3c**), consistent with the 30% FDR cut-off used for the search (refer to ‘ **Methods**’ section for more details). FDR in PCprophet is calculated by comparing hits from the provided database, in this instance CORUM, against positively predicted complexes, thereby 30% of the PPIs detected cannot be derived from CORUM. We speculated that this set of PPIs without database evidence would be localized in cellular niches with poorly characterized complexes, such as membrane bounded organelles^8^. Consistent with this hypothesis, among the top 10 most enriched GO Cellular Compartments (CC) terms for the novel PPIs, we observed localization enrichment in mitochondrion, ficolin, cytoskeleton and cytoplasmic-associated lumen (**Fig. 3d**) all with an adjusted *p*-value of less than 1%.

We identified several cases where a novel subunit is assigned to a known complex by PCprophet. For instance, PCprophet identified a novel protein complex containing the ubiquitin receptor ADRM1 and 26S proteasome (**Fig. 3e**). This association has not been reported in the CORUM database for *homo sapiens* but it has been identified in mammalian cells^21^ and is consistent with the crystal structure of *S. cerevisiae* (PDB ID: 6J2C and 6J2Q)^22^. ADRM1 is reported to be a component of the 19S proteasomal subunit in yeast^22^; accordingly we observed about 15% of the ADRM1 signal to be associated with 19S (**Fig. 3e**) while the majority was associated with the 26S proteasome, suggesting that ADRM1 is preassembled in the 19S rather than being later recruited to a fully assembled 26S. We further identified an interaction between the NEDD8 activating complex NAE1-UBA3 and ASB6 (**Fig. 3f**). The NAE1-UBA3 complex is required for cell cycle progression by transferring activated NEDD8 to UBE2M and subsequent proteasomal degradation^23^. ASB6, which belongs to the Ankyrin repeat and SOCS box (ASB) protein family, has been shown to interact with CUL5 and RBX2 to form a non-canonical E3 ubiquitin ligase complex^24^. We observed almost perfect co-elution between the NAE1-UBA3 complex and ASB6, consistent with reports of ASB6 and UBA3 co-purification in other species^25, 26^, but not with reported ASB6 binders such as CUL5^24^ (**Supplementary Fig. S3**) or reported NAE1-UBA3 binders like UBE2M^16^ or TP53BP2^16^. Taken together, the recall of protein-protein interactions absent from the training set as well as reported complexes, suggests that PCprophet can predict protein complexes in cellular models that are poorly characterized with respect to protein complexes and PPIs.

### Differential analysis of mitosis-associated protein complexes

We have identified 900 previously reported and 532 novel complexes in HeLa cell lysates derived from interphase and mitotic cells, with a similar number of complexes in each cellular state (**Fig. 4a**). Due to the continuous nature of coFrac-MS data it is possible to identify several types of profile differences at both protein and complex level. First, difference in assembly state causes a shift on the molecular mass scale, while changes at the abundance level results in an increase peak area for a particular protein. By similarity, complex compositional changes can be inferred by the difference in peak position of the subunits or addition of novel proteins, while stoichiometric changes are dependent on ratios between different proteins. This is a non-trivial issue as metrics based on profile correlation will fail in capturing abundance difference, while methods based on peak position will not detect variation in peak area. To overcome this issue, we developed a Bayesian approach to identify altered protein profiles in the different conditions tested and defined a likelihood for each interaction, which we then combine into a complex-specific likelihood (see ‘**Methods**’ for more details). This approach has several advantages over previous methods such as fold change^17^ as it does not require a pre-selected threshold and penalizes proteins with high variability. Overall, we detected 1518 proteins (238 complexes) with a probability greater than 0.5 of being differentially regulated across the cell cycle (**Fig. 4b**). On this set of proteins, we performed an enrichment analysis using GO ontology to evaluate if terms associated with cell cycle and mitosis (**Fig. 4c**) were enriched. Indeed, terms such as M phase, mitosis and nuclear division are enriched with p<0.001%. Surprisingly, we identified the Prmt5-Wdr77 complex as altered between interphase and mitosis (**Fig 4d**). This complex is composed by a hetero tetramer formed by Prmt5-Wdr77 dimers in a 1:1 ratio^27^ (PDB ID: 4GQB). While this putative 1:1 stoichiometric ratio is reflected in the protein MS intensities during interphase, upon mitosis it is significantly shifted towards a 1.75:1 ratio, indicating a gain of Prmt5 copies in the assembly relative to the composition in interphase (**Fig 4e)**. Interestingly, Prtm5^28^ and Wdr77^29^ have been independently linked to cell cycle regulation and complex stoichiometry is necessary for correct target methylation by Prmt5^27^. Thus, our data suggests a potential role for the Prmt5-Wdr77 complex in cell cycle regulation. Furthermore, key events such as activation of the master mitotic kinase complex CDK1/CCNB1 (**Fig. 4f**), increase in cohesin complex (**Fig. 4g**) and rewiring of the anaphase promoting complex/cyclosome (**Fig. 4h**) were successfully captured by our analysis strategy.

**Fig. 4.**
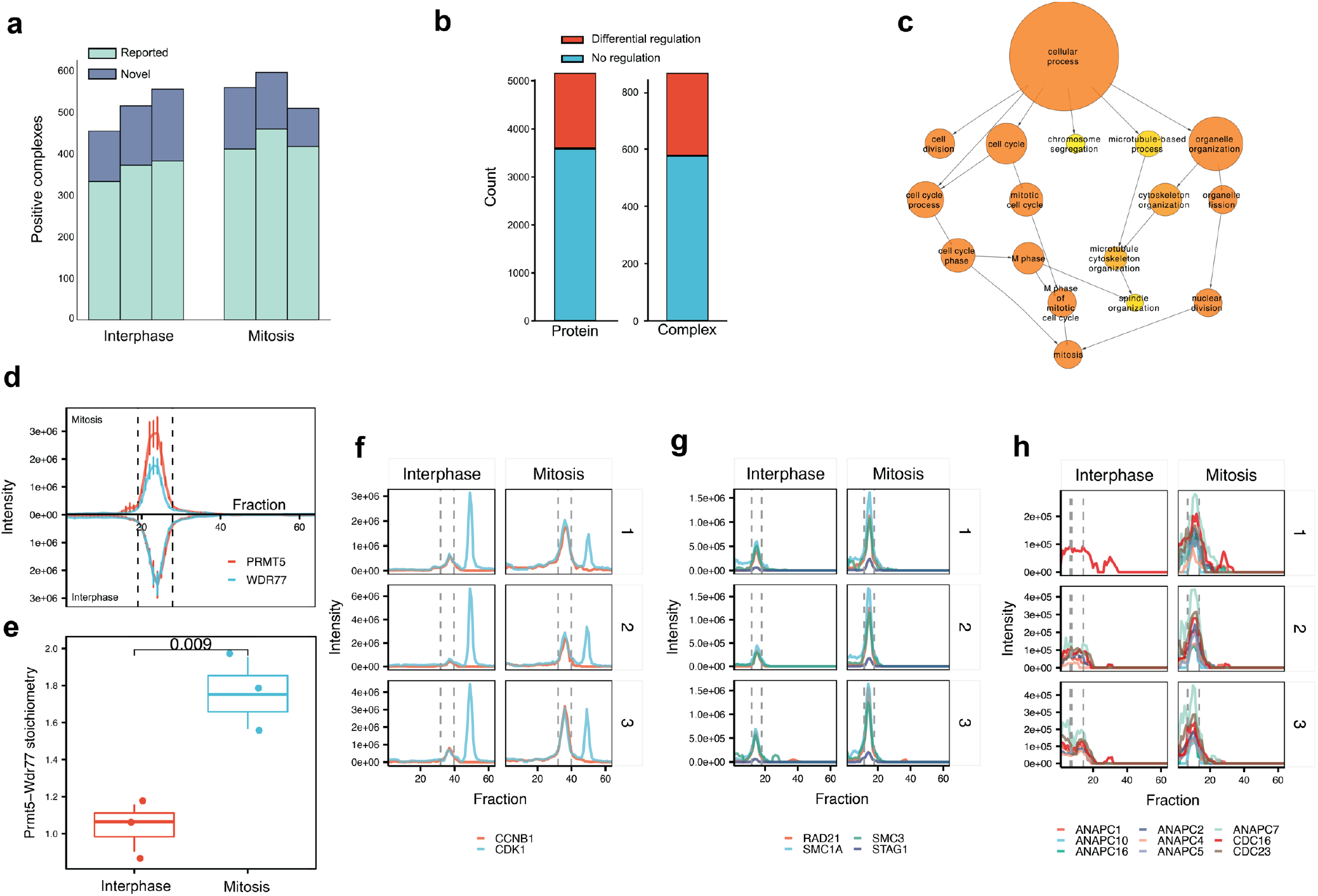
Differential analysis of complexes across the cell cycle states tested. **a**, Absolute number of novel and reported complexes in the indicated cell cycle stages. **b,** Stacked histogram for differentially regulated proteins (n=1518) and differentially regulated complexes (n=238). **c,** Enrichment for GO Biological Process for differentially regulated proteins between the two conditions using as background all proteins identified. Node size represents number of proteins within the particular category. Nodes colour represents Bonferroni adjusted *p* value, ranging from *p*=10^-3^ (yellow) to *p*=10^-8^ (orange). **d,** Mirror plot for co-elution profiles of Prmt5-Wdr77 complex in mitosis (upper positive Y axis) and interphase (negative y axis). Values were averaged across the three replicates for each condition and bar represents standard error of the mean. **e,** Bar-plot of Prmt5/Wdr77 complex stoichiometry in interphase (red, mean=1.04) and mitosis (blue, mean=1.75). **f,** Co-elution profile for the CCNB1/CKD1 complex (**G**) Co-elution profile for the Anaphase promoting complex. **h,** Co-elution profile for the cohesion complex.

To conclude, our analysis demonstrates that (i) its ability to recall known complex remodelling events in cell cycle progression, and (ii) its sensitivity to discriminate between different scenarios such as increase in abundance (**Fig. 4fg**) and difference in peak shapes (**Fig. 4h)**. Altogether, our analysis recapitulates previous knowledge about cell cycle and cell cycle-related events, selectively recalling complexes involved in cell cycle progression and mitosis.

## Discussion

Protein complexes play fundamentally important roles in mediating and regulating biological functions. Recent advances in proteomic technologies based on co-fractionation and mass spectrometric correlation profiling of protein elution patterns have opened up a promising avenue to characterize protein complexes at breadth and temporal resolution. State-of-the-art workflows such as SECS-WATH-MS techniques and complex-centric data analysis have advanced the selectivity and throughput of chromatographic protein complex detection but remain limited to the detection of previously observed protein complexes. Methods to predict novel protein complexes from co-fractionation data are based on identification of PPIs and inference of complexes from the resulting weighted network. Such probabilistic methods for network partitioning rely heavily on network topology, which makes it challenging to partition detected PPIs into complexes, due to the high dimensionality of the data. In light of this, we introduced the PCprophet framework which combines complex-level scoring with powerful machine learning technology to classify and confidently predict novel protein complexes from protein coFrac-MS data. In addition, PCprophet facilitates the differential tracking and comparison of these complexes across two or more experimental conditions that become increasingly accessible via high throughput implementations of coFrac-MS. We have demonstrated outstanding prediction performance of PCprophet on manually annotated datasets and have shown that the method significantly outperforms state-of-the-art complex prediction and identification tools. We have developed a Bayesian inference-based method to analyse differences in protein complex abundance and composition across conditions. Our analysis on proteomic profiles across the cell cycle of HeLa cells demonstrated that PCprophet can capture expected changes in protein complexes between interphase and mitotic cells.

PCprophet is available in command-line version under MIT licence (https://github.com/fossatiA/PCprophet) and is easily applied to any coFrac-MS dataset. The data pre-processing module readily accepts different types of quantitative protein level tables. PCprophet could also be applied in clinical proteomics and personalized medicine areas, to assist the discovery and analysis of novel protein complexes and to identify complexes that are altered across groups of samples. We anticipate that the PCprophet package will serve as a reliable and accurate tool for novel protein complex prediction and analysis from co-fractionation MS data because it extends the scope of comparative proteomics from the level of differentially abundant proteins to the level of differentially abundant and perturbed complexes between samples, thus bringing proteomic analysis closer to biological function.

## Methods

Methods, including statements of data availability and any source code and references, are available in the online version of the paper.

## Acknowledgements

This work was supported in part by the Swiss National Science Foundation (grant No. 3100A0-688 107679 to R.A.) and the European Research Council (ERC-20140AdG 670821 to R.A.). C.L. is currently supported by a National Health and Medicine Research Council (NHMRC) of Australia CJ Martin Early Career Research Fellowship (1143366). M.G. and F.F. acknowledge the support by the Innovative Medicines Initiative project ULTRA-DD (FP07/2007-2013, grant No. 115766). I.B. acknowledges funding support from the Swiss National Science Foundation (31003A_166435). AWP is supported by a NHMRC Principal Research Fellowship (1137739). The authors would like to thank Dr. Natalie de Souza from ETH Zurich and Associate Professor Jiangning Song from Monash University for their critical comments on this study.

## Author contributions

R.A., A.F., C.L. and M.G. conceived and designed the project. A.F. and C.L. designed, developed and implemented PCprophet, and conducted data analysis, machine-learning prediction and benchmarking experiments with other existing methods. P.S. designed and implemented the differential analysis module for protein complexes. M. Heusel, F.F. and F.U. contributed to data annotation and provided critical feedback and comments on the biological aspects. M. Hallal contributed to EPIC performance comparison and reproducibility test for PCprophet performance. I.B. assisted with benchmarking experiments with CCprofiler and provided useful insights. C.T.K., P.X. and A.W.P. provided critical and insightful comments during the development of PCprophet. C.L., A.F., M.G. and R.A. drafted the manuscript, which has been revised and approved by all the other authors.

## Competing financial interests

The authors declare no competing financial interest.

## Methods

### Training dataset curation and annotation

In total, three co-fractionation replicates using SWATH and DDA-SILAC based datasets were used to train and evaluate PCprophet, including the SEC-SWATH-MS dataset from HEK293 cell line^10^ for training PCprophet, the mitotic proteomic data from HeLa CCL2 cells^17^ and the DDA-SILAC dataset extracted from the study by Kristensen and used as testing dataset for PrInCE^12^, for independently testing PCprophet. Note that we did not use the *C. elegans* protein complex dataset from the EPIC^8^ package to test PCprophet for the following reasons: (i) their datasets used spectral count; however our previous study showed lower performance of spectral counts compared to XIC based quantitation (MS1 or MS2) for complex analysis^10^; (ii) the features and pre-processing employed in PCprophet are inherently continuous in nature such as correlation and FWHM; and (iii) EPIC was developed and evaluated using the same dataset. It is therefore challenging to conduct an unbiased and fair estimation and comparison of the performance for all the other approaches. A structuralized description of these datasets is available in **Supplementary Table S2**.

To train accurate machine-learning models, we manually annotated the protein complexes from the SEC-SWATH-MS dataset from HEK293 cells^10^. Briefly, samples were acquired in SWATH mode using a sample-specific library generated from high-pH fractionated samples. Following conversion to mzXML via msConvert, OpenSWATH search was performed with the parameter previously described^10^. As a result, the final feature alignment outputs from TRIC^30^ (using top2 protein quantification) was used for the training PCprophet. The reference core (non-redundant) complexes were downloaded from CORUM v3.0^13^. Protein accession numbers were converted into gene names and sequentially mapped to CORUM. We removed complexes from our dataset where the number of subunits present in the dataset was less than 50% of known components to retain only complexes with high coverage, consistent with the annotation strategy for the complex analysis software CCprofiler^10^. To train a supervised classification model, it is crucial to reliably annotate the samples of positive (i.e. complexes with good co-elution profiles) and negative (i.e. complexes with poor co-elution profiles) classes. For the training dataset (*i.e*. HEK293), a protein complex was annotated positive if it satisfied the following criteria. First, more than 75% of the known subunits coeluted in the same fraction with baseline resolution; second, the main peak was required to have a minimal FWHM (full width at half maximum) of 4 fraction; third, minimal normalized height of 20% to the maximum signal for every protein and needs to be at least 10% above background, and the complex has been annotated in the CORUM database. On the other hand, if the complex was annotated in CORUM but did not pass all the other criteria it was annotated as negative. To objectively annotate the protein complexes in the training dataset, three annotators were involved in this procedure and only positive and negative protein complexes nominated and agreed by all the annotators were used. Notice that we did not randomly select proteins to form negative but ‘fake’ protein complexes, as there would be a huge number of different combinations and possibilities and might result in random selection of undiscovered protein-protein interaction. In addition, we changed the original number of fractions of the training dataset from 81 to 72, to standardize number of fractions across different experiments. The final resulting training dataset contained 242 positive protein complexes and 738 negative complexes.

### Dataset pre-processing

Prior to the generation of potential protein complexes based on the protein raw matrices across various conditions, four data pre-processing steps are performed, including Gaussian filtering, missing value imputation, linear interpolation, and data rescaling. One-dimension Gaussian filtering,

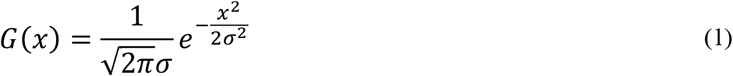

was employed to smooth the data by removing noise and approximating peaks as Gaussian curves, where *x* denotes the intensity of the current fraction and *σ* was set to 1. To remove the missing values in the raw data, an imputation strategy by calculating the average of the two neighbours of missing value, was implemented. The number of fractions *N* in the co-fractionation experiments always varies due to various experimental setups and inherent variability. When constructing the machine-learning models, the number of features is dependent on the number of fractions (see ‘Feature engineering and construction of machine-learning models’ for feature generation). Based on the number of fractions (i.e. 72) in our training dataset (see ‘Data curation and annotation’ for details), it was therefore necessary to rescale the number of fractions of user-provided datasets to 72. In PCprophet, resampling and one-dimensional linear interpolation were applied for this purpose, thereby rescaling the number of fractions to 72, consistent with the training dataset. Lastly, we added an additional step to standardize every protein profile from their original intensity in the range [0, 1] to make it independent from the quantitation strategy used.

### Hypothesis generation

In this study, hypothesis generation refers to the construction of putative protein complexes. Theoretically, there could be a huge number of possible combinations of proteins to form different protein complexes. Rather, we proposed a hypothesis generation module to construct potential protein complexes by aligning peaks of different proteins and cutting the dendrogram-like tree structure, similar to the procedure discussed in a previous study^11^. To do so, hypothesis generation firstly performs peak-picking to identify all the apexes of intensity and their associated fractions of all proteins in the input data, with the help of the Python package ‘SciPy’ (https://www.scipy.org/). Then linkage hierarchical clustering using the ‘Ward’ distance measure was performed based on the apexes collected during the peak-picking stage. A dendrogram-like tree structure was then generated based on the results of the linkage hierarchical clustering. The hypotheses (i.e. putative protein complexes) were then generated by cutting the dendrogram from bottom to top at each level. A conceptual illustration of the hypothesis generation is presented in **Supplementary Fig. S4**. In practice, these steps are performed on a linkage matrix instead of a dendrogram structure to further reduce the computational burden.

### Feature engineering and construction of machine-learning models

To represent a protein complex, we designed a variety of features based on the protein co-elution profile and the number of fractions *N*. These features are mainly categorised in four groups: (1) average intensity difference of proteins within each fraction (**Supplementary Fig. S5a**), (2) local correlation of proteins at each window (**Supplementary Fig. S5b**), (3) shift of apex fraction of each protein (**Supplementary Fig. S5c**), and (4) average full width half maximum (**Supplementary Fig. S5d**).

For the intensity difference of proteins and the correlation of proteins at each fraction, we set a sliding window (6 fractions wide) with 1 fraction step wise increase. The intensity difference of proteins, *D_intensity_i_* reflects the average difference of intensity values of protein *a, b*, and *c*, at each fraction, calculated by:

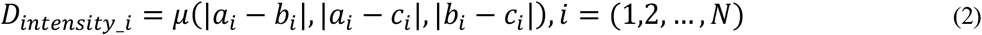

where *i* denotes the number of fractions. As a result, the dimension of this feature type is *N*. Similarly, at each window, pairwise correlation of intensity was also calculated, using:

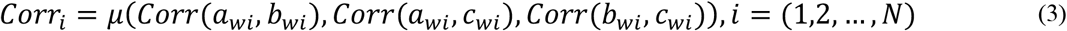

Where *a_wi_* denotes the local value of *a* in a window *w* centred at fraction *i*. For the other two types of features, including fraction difference of apex peaks and full width half maximum a two step-procedure is employed. First, all peaks for a protein complex hypothesis are selected, and then a modified version of the Dijkstra’s algorithm is applied to select the peaks for every protein with the minimum distance. By selecting the closer peaks, we are able to positively predict proteins with multiple peaks in separate assemblies, as we avoid the use of heuristic to select the complex-specific peak. While the apex difference is a feature used also in the PrInCE software^12^, substantial differences are present as in this tool, the fraction with the maximum value for every protein is counted as apex, thereby using always the same peak for a protein in multiple assemblies.

For the average apex difference, we used following formula for the calculation, respectively:

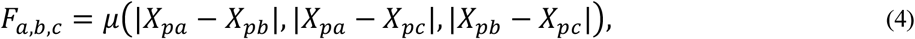

where *X_pa_, X_pb_, X_pc_* represent the apex fraction of protein *a, b* and *c*, respectively. The average full width half maximum is calculated using:

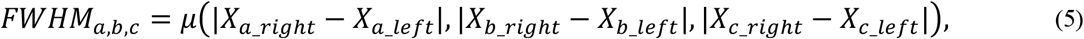

while *X_a_right_*-*X_a_left_, X_b_right_, X_b_left_, X_c_right_, X_c_left_* demonstrate the width when achieving half intensity area of the co-elution curve of protein *a, b* and *c*, respectively. In total, we generated *2N* + 2 features (i.e., 146 when *N*=72).

Five well-established machine-learning models were selected to test the prediction performance, including Decision Tree (J48)^31^, Random Forest^32^ (RF), Naïve Bayes^33^ (NB), Support Vector Machines^34^ (SVM) and Logistic Regression^35^ (LR). For SVM, we selected two major kernels, including polynomial and RBF^36^ (Radial Basis Function) kernels, due to the consideration of the balance of computational complexity and running time. These two models were then termed as SVM_POLY and SVM_RBF, respectively. Note that the above machine-learning algorithms were implemented using the scikit-learn package^37^ and cross-tested in the WEKA^38^ platform. Different implementations of such algorithms in other platforms may cause difference in term of prediction performance. To objectively portrait the prediction performance and avoid overfitting, five-fold cross-validation strategy using the training dataset was performed, together with five widely acknowledged performance measures, including accuracy (ACC), area under the curve (AUC), Matthew’s correlation coefficient (MCC), sensitivity and specificity:

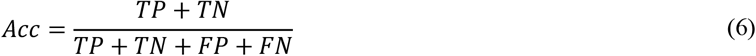

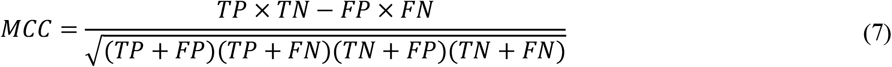

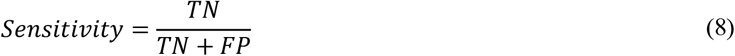

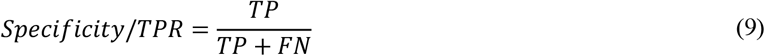

where *TP, TN, FP, FN* are true positives, true negatives, false positives and false negatives, respectively.

### Benchmarking with state-of-the-art approaches for co-fractionation MS based protein complex prediction

We compared the prediction performance of PCprophet with currently existing computational approaches for protein complex characterization based on co-fractionation MS data, including CCprofiler^10^, EPIC^8^, and PrInCE^12^. CCprofiler is a statistical approach for the identification of protein complexes by referencing databases as prior information. In contrast, EPIC and PrInCE were designed to infer protein complexes based on PPI prediction from co-elution profiles. When running EPIC, two provided models, including SVM and RF (i.e. -M RF and -M SVM) were both tested with other parameters by default (i.e. -t 9606 for *H. sapiens;* -s 11101001; -f STRING). Two datasets, as shown in **Supplementary Table S2** were used to benchmark with CCprofiler and EPIC, including the mitotic proteomic data from HeLa cells^17^, and the soluble protein complex dataset from the study of Stacey *et al*.^12^

#### Benchmarking using HeLa mitotic proteomic data

The TRIC feature aligned file of the dataset was imported into CCprofiler and following sibling peptide correlation (the ‘filterBySibPepCorr’ function). Protein quantification was done using the top 2 proteotypic peptides per protein. The resulting protein tables were exported and used as input for EPIC and PCprophet. PCprophet was run with default parameters, with the FDR fixed at 30% and controlled using the CORUM database. CCprofiler complex-centric analysis was done as previously described^1^, using smoothing_length = 9, corr_cutoff = 0.95, window_size = 8, rt_height = 3 and a 2x molecular weight cutoff^1^. For the calculation of recall against CORUM, PCprophet output (i.e. the ‘ComplexReport.txt’ file) was filtered to only ‘Reported’ complexes which were predicted as ‘Positive’; while EPIC derived complexes were matched to CORUM by defining a positive predicted complex in which 50% or more subunits are reported in a single CORUM core complex. For CCprofiler, the positive complexes were defined when the q-value is smaller than 5%, as previously described^1^. The number of complexes was considered across replicates as the number of unique positive CORUM Complex ID. Two measures were applied when assessing the prediction performance, including True Positive Rate (TPR; specificity) and Positive Predicted Value (PPV), which is defined as follows:

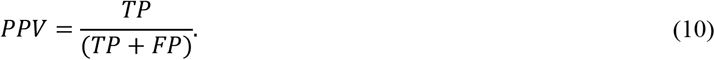

For evaluation of average network degree, protein complexes from the prediction outputs of EPIC and PCprophet, and CORUM (Human only) were collapsed to a PPI network. The reference networks from STRING (Human) and BioPlex were downloaded and used directly. Degree calculation for every protein in the networks was done using the NetworkX package v2.1 (https://networkx.github.io) and ranked. A log-log plot was generated and the AUC for the resulting curve was calculated using the integrate module from SciPy (https://www.scipy.org). For evaluation of node centrality, the resulting complexes from PCprophet, EPIC_RF and EPIC_SVM were projected into a subgraph generated by filtering STRING to only nodes present in the original protein matrixes thereby representing all the reported protein-protein interaction available in our data. For every complex in the three tools (PCprophet, EPIC_RF and EPIC_SVM) the average complex closeness (ACC) was defined as the mean shortest path between all members. The resulting vector represents the tendency of the different algorithms to recapitulate first, second or outer shells level of interactors. The same was done also for CORUM complexes within the same subgraph. Average complex size was defined as the mean number of subunits for the same complex across replicate for PCprophet and the number of subunits for every complex in the EPIC output.

#### Benchmarking using the DDA-SILAC dataset^18^

Condition1.tsv and condition2.tsv were downloaded from https://github.com/fosterlab/PrInCE-Matlab and separated into the different replicates. PCprophet and EPIC were run as described above. TPR and PPV were calculated as described above for CORUM. For BioPlex and STRING both networks were filtered to remove proteins not identified. PCprophet, EPIC_RF and EPIC_SVM derived complexes were collapsed to generate a PPIs network and then recall was calculated using a PPI-centric approach by assessing which fraction of PPIs was present over the entire reference (TPR) and which fractions of PPIs was correct across all of the predicted one (PPV). Centrality assessment via KS test and AUC calculation was done as described above.

### Post-prediction processing

During this stage, GO (Gene Ontology) term score filtering and complex combination and collapsing are performed in order to ensure the reliability of predicted complexes. Despite the incompleteness of GO term annotation of CORUM database, we compared the distributions of GO terms of predicted protein complexes and documented protein complexes in the CORUM database^13^ (**Supplementary Fig. S6a**). We first collected GO terms of each protein in a predicted complex based on the annotation of AmiGO2 database (i.e. the Gene Ontology resource)^39–41^. Then, for every possible protein pair we calculated pairwise GO term semantic similarity using the strategy published by Wang *et al*^42^. For instance, given a protein complex *PC* with three subunits *A, B* and *C*, all the GO terms, including molecular function (MF), biological process (BP), and cellular component (CC) are collected. For all the three possible protein pairs, including *A-B, A-C* and *B-C*, within each category, the semantic similarity scores of all the pairwise GO terms are calculated and the average score is reported as the overall score for the current GO category. The final overall GO score of protein complex *PC* in this case is then defined as follows:

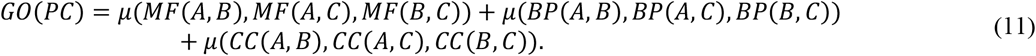

The GO term scores of core protein complexes from the CORUM database are calculated using the same strategy. Given the two distributions of the GO term scores of both protein complex hypothesis and the complexes harboured from the CORUM database, we then estimated false discovery rate for the positively predicted hypothesis by calculating global FDR for every GO scores of positive CORUM complexes with the following formula:

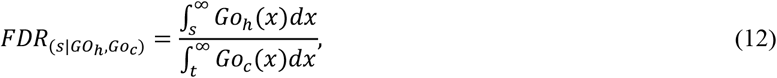

where *Go_h_* and *Go_c_* are defined as distributions of the Wang similaritydifferential regulation of protein abundance score for hypothesis (h) and CORUM-derived complexes (*c*). This allows us to obtain the specific GO term score that satisfies the target FDR for filtering the predicted protein complexes without having to use a fixed threshold, thereby allowing for more or less conservative searches.

Given the possibility that the positive complex hypothesis might be a subset of a bigger complex or might contain multiple smaller complexes, PCprophet allows users to select a complex collapsing mode to further process the predicted complexes (**Supplementary Fig. S6b**), including ‘**GO**’ (based on GO terms), ‘**CAL**’ (based on the provided calibration curve), ‘**SUPER**’ (to find the biggest protein complex), and ‘**NONE**’ (to ignore this process). Specifically, following prediction and FDR control, overlap is calculated and complexes for which the overlap is more than 0.75 are merged on the different criterion. Given a set of complexes defined as follows in **Supplementary Table S3**, for example, collapsing using ‘**GO**’ will results in the complex which has the greatest GO score (i.e. **PC2**). The ‘**SUPER**’ option will select the of the complex with the highest number of subunits (i.e. **PC3**) while choosing ‘**CAL**’ will calculate the difference between apparent MW from the SEC and extrapolated molecular weight from the calibration curve. The complex with the smaller difference (i.e. **PC3**) will therefore be selected. The ‘**CAL**’ mode is selectable only if the calibration and a molecular weight table from the UniProt^43^ database or similar format is provided. ‘**NONE**’ option will skip the collapsing procedure.

### Protein complex differential analysis across different conditions

#### Bayesian inference of differential regulation of protein abundance

Inferring differentially regulated proteins assumes that protein abundance measurements were obtained for a number of samples which differ in a biological phenotype of interest. This situation allows representing the protein abundance measurements for one protein as a matrix ***X*** where rows correspond to samples and columns correspond to retention time. Phenotype information ***t*** is assumed to be discrete with cardinality *#**t*** and order matched such that the phenotype information for sample *n, t_n_ =* t[n] corresponds to the protein abundance row vector *x_n_* = ***X***[*n*]. If we assume that the correct model is among the investigated candidate models, we have a problem termed ‘m-closed model selection’^44^. In this situation the Bayesian approach to inferring whether a phenotype change corresponds to differential regulation in protein abundance suggests to use marginal likelihoods to derive the corresponding Bayes factors^44^. To obtain a solution which may be calculated analytically, we use the model illustrated in **Supplementary Fig. S7a**. The model represents protein abundance measurements by Normal-Whishart distributions. Differential regulation is implicitly represented (variable not shown in the graph) via a protein specific indicator variable *I_p_*. Differential regulation is coded by *I_p_* = 1 and results in modelling the protein abundances *x_n_* conditionally on phenotype states *t_n_* by phenotype specific Gaussian distributions. To obtain a measure of differential protein regulation we compare the *I_p_* = 1 model with a simpler explanation which we denote as *I_p_* = 0. The simple model corresponds to a non-differential regulation assumption and uses one common Gaussian distribution to model *x_n_* irrespective of the phenotype state *t_n_*. Irrespective whether we have *#t* multivariate Gaussians in case of *I_p_* = 1 or one shared Gaussian in case of *I_p_ = 0*, the joint distribution of data and model parameters *P*(***X**, μ,Λ|**t**,γ,m, g,h*) is represented by the directed acyclic graph (DAG) in **Supplementary Fig. S7a**. In case of *I_p_* = 1 we have

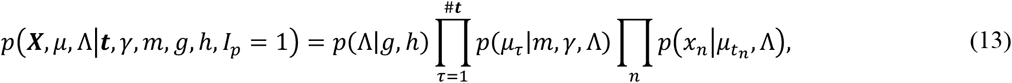

whereas in case of *I_p_* = 0 we have the simpler relation

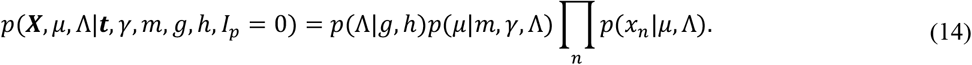

The next step to obtain a measure of differential regulation is to calculate the marginal likelihood for both models in Equation (13) and Equation (14). For Equation (13) we get

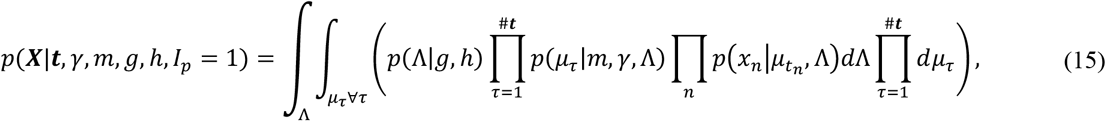

while Equation (14) leads to

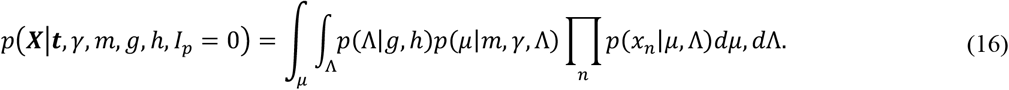

For coding equal preference for the indicator values *I_p_* = 1 and *I_p_* = 0 we use a flat prior and hence *P(l_p_* = 1= = *P*(*I_p_* = 0) = *P*(*I_p_* = 0) = 0.5. The marginal likelihoods in Equations (15) and (16) can subsequently be converted to the posterior probability for differential regulation of protein abundance *P*(*I_p_ ≡ 1|**t**,γ,m,g,h*):

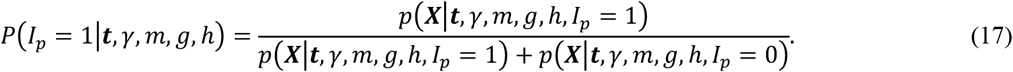

Taking the posterior probability *P*(*I_p_ ≡ 1|**t**,γ,m,g,h*) in Equation (17) as measure of differential protein regulation is justified by the fact that Bayesian model selection has Occam’s razor built in^44^. Posterior probability values *P*(*I_p_ ≡ 1|**t**,γ,m,g,h*) which are larger than 0.5 will only be observed if the more complex model (*I_p_* = 1) provides a substantially better fit of the data ***X*** and ***t*** than the simpler model (*I_p_* = 0).

#### Inferring differentially regulated protein complexes

Inference of differential regulation of protein complexes assumes that the assignment of proteins to protein complexes is known. All subsequent derivations assume thus that the protein complex *c* is defined as a set of proteins *C_c_* = [*P_1_,P_2_,…,P_c_*] of cardinality *C*. We assume furthermore that a set of retention profiles *X_c_* = [*X_P_1__,X_p_2__,…,X_p_c__*] and a corresponding set of phenotype descriptions *t_c_* = [*t_P_1__, t_p_2__, …, t_P_c__*] is available. If the retention profiles in *X_c_* and thus the corresponding phenotype characteristics in *t_c_* can at least in part be paired among all proteins which establish a complex, we have the subset *N_c_* = [*n*_1_, *n*_2_, …, *n_K_*] of samples for which complete observations are available. To prepare inferring differentially regulated protein complexes we may in this situation aggregate the protein specific retention profiles to a column concatenated matrix ***Y**_c_* which represents all retention profiles of the entire complex. Denoting the selection of the n^th^ row of matrix *X_P_c__* as *X_P_c__*[*n*] and column wise row concatenation of row vectors *X_p_c__* [*n*] and *X_p_c+1__*[*n*] as [*X_p_c__*[*n*], [*X_p_c+1__*[*n*]], we obtain

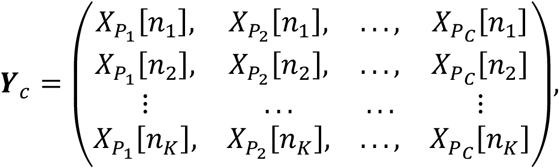

as protein complex specific matrix of retention profiles and ***u**_c_* = [*t_p_1__* [*n_1_*],*t_p_1__* [*n_2_*],…,[*t_p_1__* [*n_K_*]]^T^ as protein complex specific phenotype vector. Inference of differentially regulated complexes is now a straightforward application of Equation (15), Equation (16) and Equation (17). We have just got to replace the protein specific retention profiles ***X*** in these equations with the protein complex specific retention profiles ***Y**c* and exchange the phenotype characterization *t* with the phenotype characterization of the protein complex *u_c_*. Pooling of retention profiles requires in addition to the assumptions which led to the DAG in Figure 1 no additional assumptions. While this is an advantage of the approach, we have to consider that pooling of samples requires compete sets of paired retention profiles which have to be available for all proteins which aggregate to the complex. In practice measurement errors will lead to random dropouts and thus to a potentially small number of samples where all data is available. To avoid such information loss by pairing of samples, we propose an additional approach for assessing differentially regulated protein complexes by Bayesian model probabilities. For assessing differential regulation of protein complex *c* we apply Equations (15), (16) and (17) for every protein *P_k_* ∈ *C_c_* separately. Following the assessment on protein level, differential expression of protein complex *c* is coded via a binary indicator variable *C_c_*. Assuming conditional independence among proteins we may represent this proposition by the DAG in **Supplementary Fig. S7b**. The DAG leads for the posterior probability of differential regulation of protein complex *C_c_* finally to Equation (18).

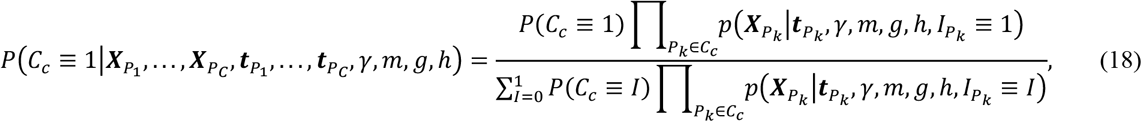

with *P_k_* ∈ *C_c_* denoting all proteins which aggregate to the protein complex *c*. The prior probability for protein complex *c* being differentially regulated, *P*(*C_c_*), is assumed to be identical for both indicator values and thus *P*(*C_c_* = 1) = *P*(*C_c_* = 0) = 0.5. The expression *p*(***x**_Pk_|**t**_Pk_,γ,m,g,h,I_Pk_* = [0,1]) denotes for *I =* [0,1] the marginal likelihoods we obtain with the model in **Supplementary Fig. S7a** for protein *P_k_* according to Supplementary Equations (15) and (16).

### Software implementation and data visualisation

The command-line version of PCprophet was implemented and visualised using Python, together with third-party packages including SciPy, Pandas^45^, scikit-learn^37^, NetworkX^46^, and Matplotlib^47^.

### Source code availability

PCprophet is open-access and freely available for academic purposes at https://github.com/fossatiA/PCprophet under the MIT License.

## Supplementary Methods

### Mathematical details for Bayesian inference of differential regulation of protein abundance and protein complexes

We now present further mathematical details how we may express the marginal likelihoods in Equations (15) and (16). We start this derivation by expressing the prior densities over *Λ* and *μ* for the simpler model *I_p_* = 0 and the densities over *μ_τ_∀_τ_* for the more complex model *I_p_* = 1. As is mentioned above, the prior over the precision matrix *Λ* is coded as a product of Gamma densities. With Λ = *diag*([*λ_1_,…,λ_D_*]) and *D* denoting the input dimension (number of columns) of ***X*** we get

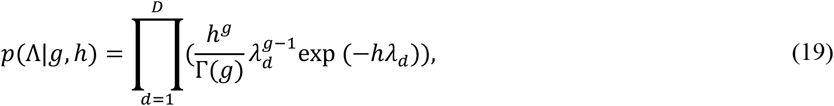

where 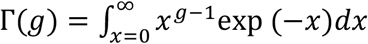 denotes the Gamma function. The multivariate Gaussian prior over μ for the non-differentially regulated case *I_p_* = 0 is

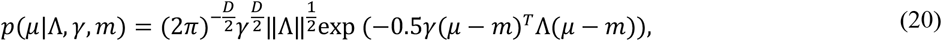

where ||Λ|| denotes the determinant of the precision matrix. Finally, we get the multivariate Gaussian prior over μ for the differentially regulated case *I_p_* = 1 as

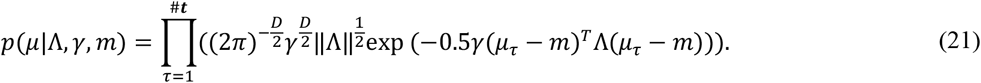

To calculate the marginal likelihood for the model *I_p_* = 1 we integrate Equation (13) first with respect to all *μτ* and then with respect to Λ to get

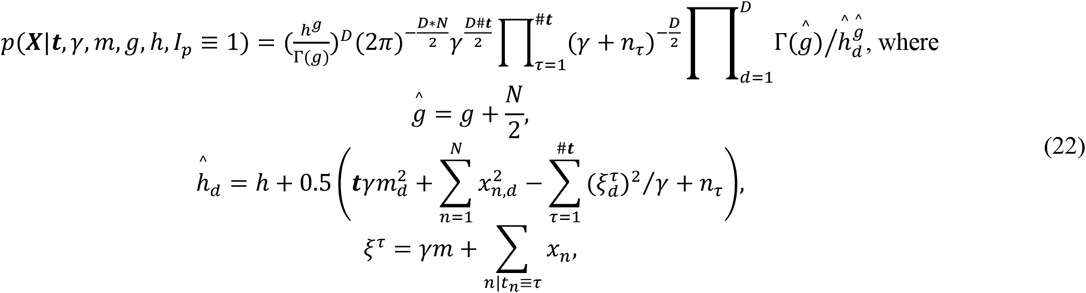

and we use *N* to denote the number of all samples and *n_τ_* to denote the number of samples which have phenotype level *τ*. The final expression for the marginal likelihood of the simpler model *I_p_* = 0 from Equation (16) is easily obtained from Equation (22). We just have to replace *nτ* with *N* and *#**t*** with 1. We should note that for numerical stability we calculate log (*p*(*X|t,γ,m,g,h,I_p_*)). The calculations reported in this paper set the hyper parameters for *g, h* and *γ* to *g* = 0.8, *h* = 1.5 and *γ* = 0.025. As prior location *m* we use the sample mean or set *m* = 0.

## Supplementary Results

### Optimization of PCprophet machine learning framework for complex prediction

In order to reach optimal performance in correctly classifying protein complex signals from the co-fractionation datasets, we explored different types of machine learning strategies and their performance to recall a set of manually curated protein complex signals in a previously published dataset^10^. To train the machine learning models, we used manually annotated data (refer to the ‘**Methods**’ section for more details) using criteria similar to the strategy applied in Heusel *et al*^10^. As the negative complexes significantly outnumbered the positive complexes (i.e. 738 *vs*. 242) based on our manual annotation, we evaluated the performance of PCprophet on two instances of the input dataset: one where all the negatives were used and the other where an equal number of negatives as positives were randomly selected We tested the performance of PCprophet based on five well-established machine learning models using five-fold cross-validation including Decision Tree (J48)^31^, Random Forest^32^ (RF), Naïve Bayes^33^ (NB), Support Vector Machines^34^ (SVM) and Logistic Regression^35^ (LR) algorithms[Figure comparing the performance of the different algorithms. We determined that the RF achieved its best performance when the number of trees was set to 500 via a separate stratified 10-fold cross-validation on the entire dataset (**Supplementary Fig. S8**); for all other algorithms, we used default parameters. In the latter dataset, we performed 100 trials for this random selection procedure (**Supplementary Table S4).** RF outperformed all the other machine-learning algorithms and selected as the algorithm for PCprophet, for example, achieving an AUC of 0.991, accuracy of 96.9% and MCC of 0.916, irrespective of the number of negative complexes used (**Supplementary Fig. 9; Supplementary Table S4**). To build a balanced and unbiased classifier, we rebuilt the RF model on the dataset with equal positive and negative complexes, on which RF achieved the best performance according to the 100 trials of 5-fold cross-validation. This rebuilt RF model is then used as the core predictor for PCprophet.

## Supplementary Figures

**Supplementary Fig. S1.**
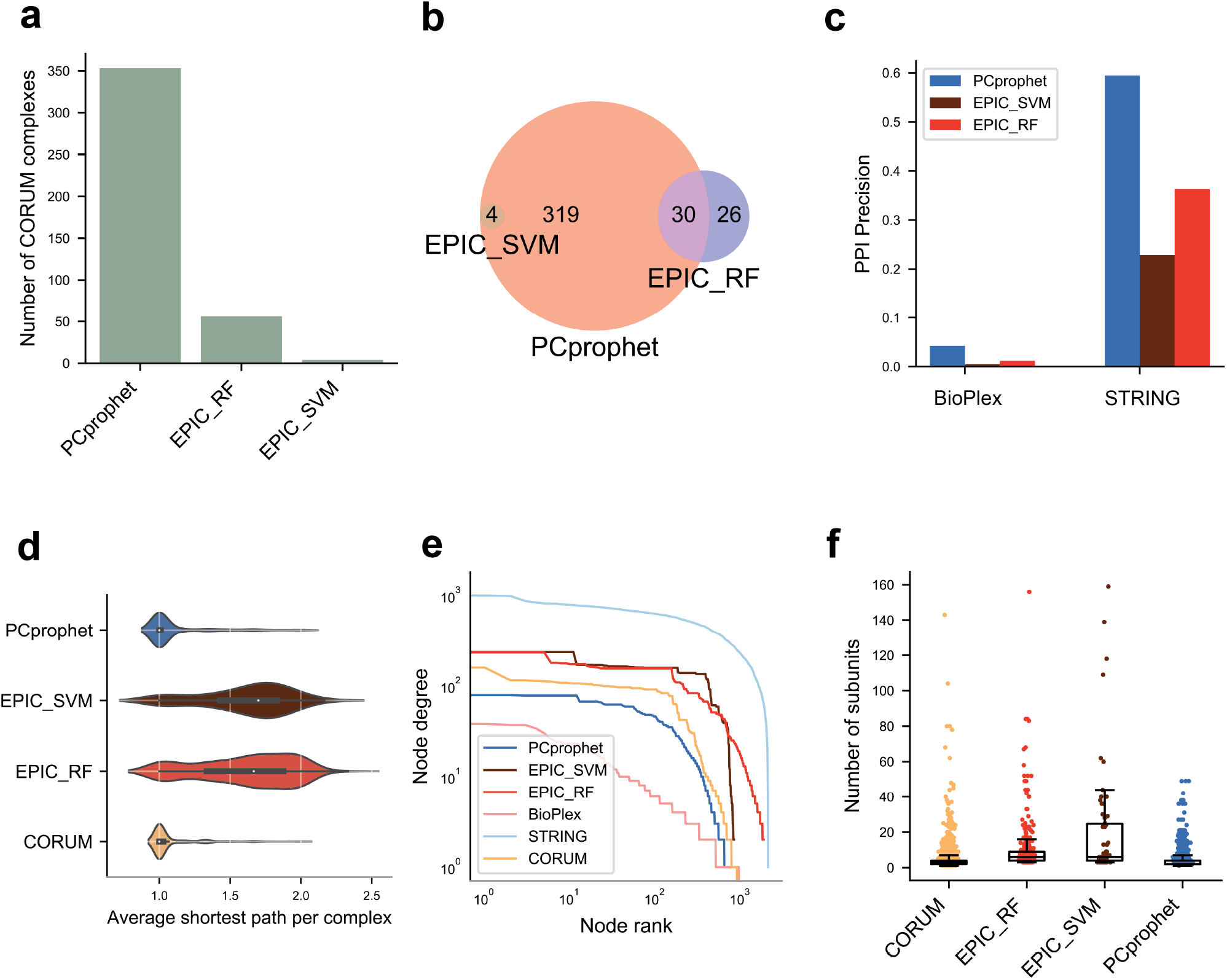
Benchmarking PCprophet with state-of-the-art software for complex profiling on the DDA-SILAC dataset. **a**, Absolute number of CORUM complexes recovered by each tool. **b**, Complex IDs overlap across all tools. **c**, Precision of PPI prediction for *de novo* protein complex prediction tools. **d**, Distribution of shortest path per complex across all subunits. The medians are highlighted using the white dots. **e**, Log-log plot showing the topology of the network generated by each tool *versus* ground-truth databases. **f**, Number of subunits per complex across different tools. Boxplot shows the medians and the ticks represent standard deviation.

**Supplementary Fig. S2.**
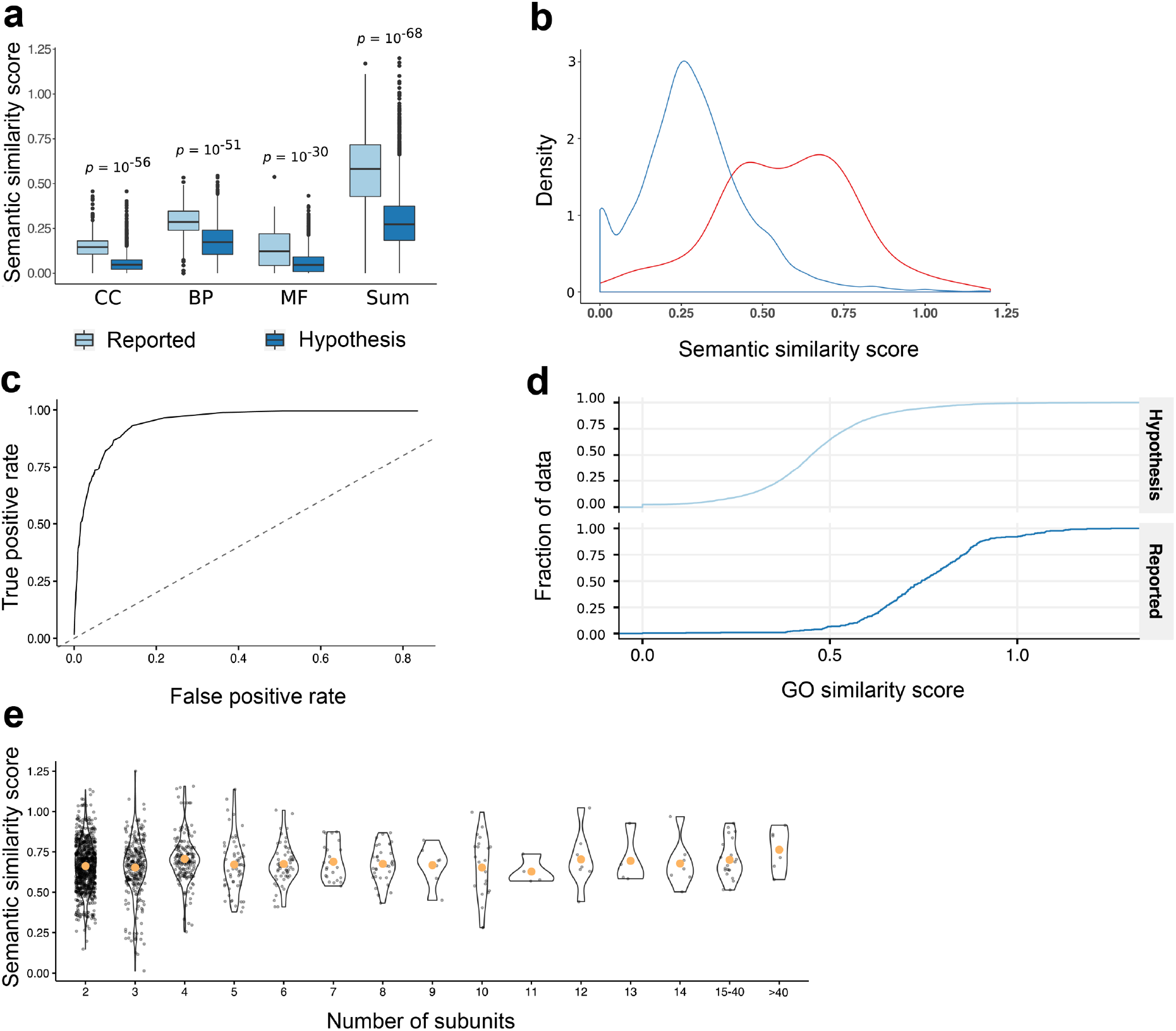
Evaluation of GO score for estimating the false discovery rate. **a**, The boxplot illustrating the separation of hypothesis and CORUM using the three individual ontologies or the combination of the three (i.e. Molecular Function – MF; Biological Process – BP; Cellular Component – CC). **b**, The density plot showing the separation of hypothesis and ground-truth CORUM database using the sum of the three ontologies. **c**, Performance of GO term for separation of true and false PPIs. **d**, Empirical cumulative distribution plot between hypothesis and reported complexes from CORUM. **e**, Distribution of the sum of the three GO ontologies across different subunits size for reported complexes.

**Supplementary Fig. S3.**
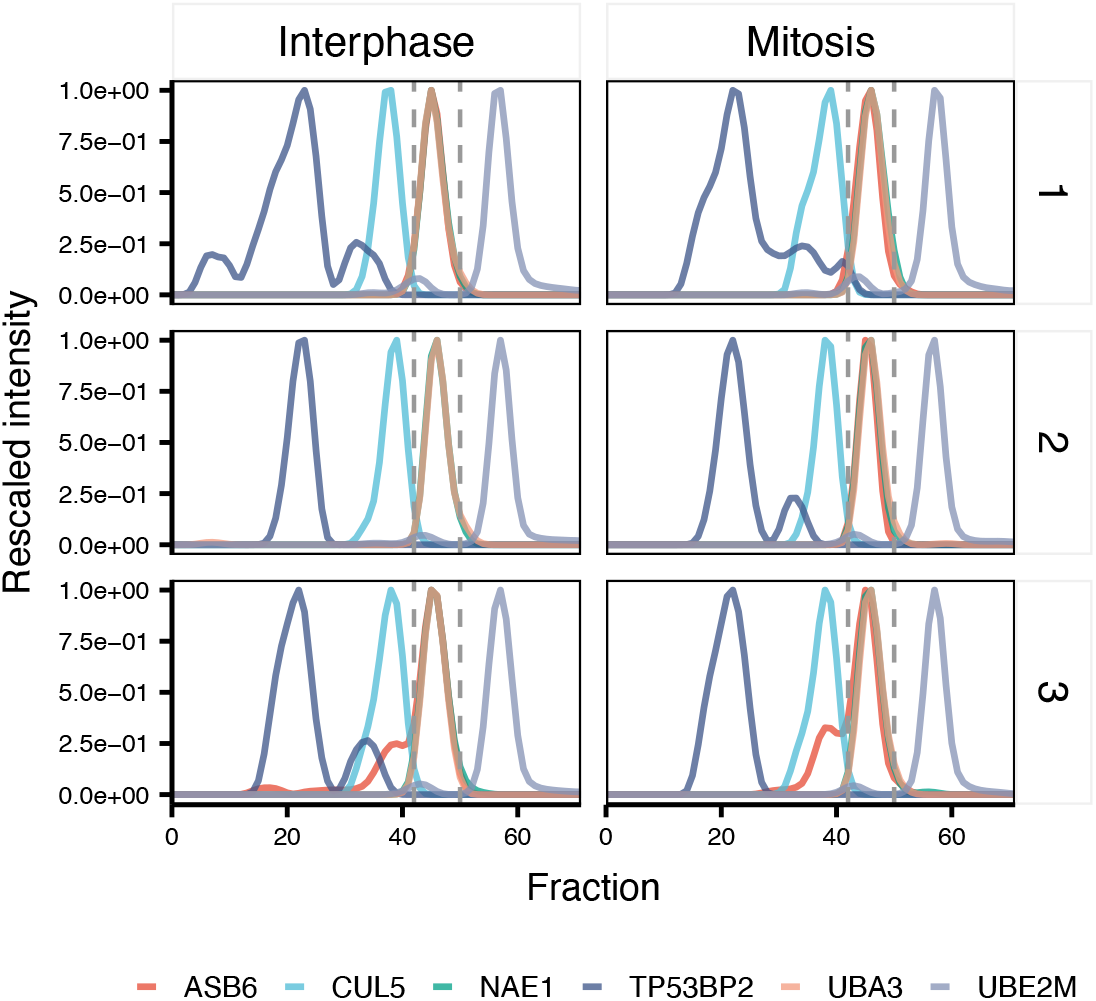
Elution profiles of reported binders for ABS6 (CUL5) and NAE1-UBA3 (TP53BP2, UBE2M) across the two experimental conditions and the three replicates. The region between the dotted lines represent the peak position of the novel ASB6-UBA3-NAE1 complex. The absence of coelution of reported interactors between the dotted lines suggests the presence of a novel complex rather than co-occurrence of complexes of similar size.

**Supplementary Fig. S4.**
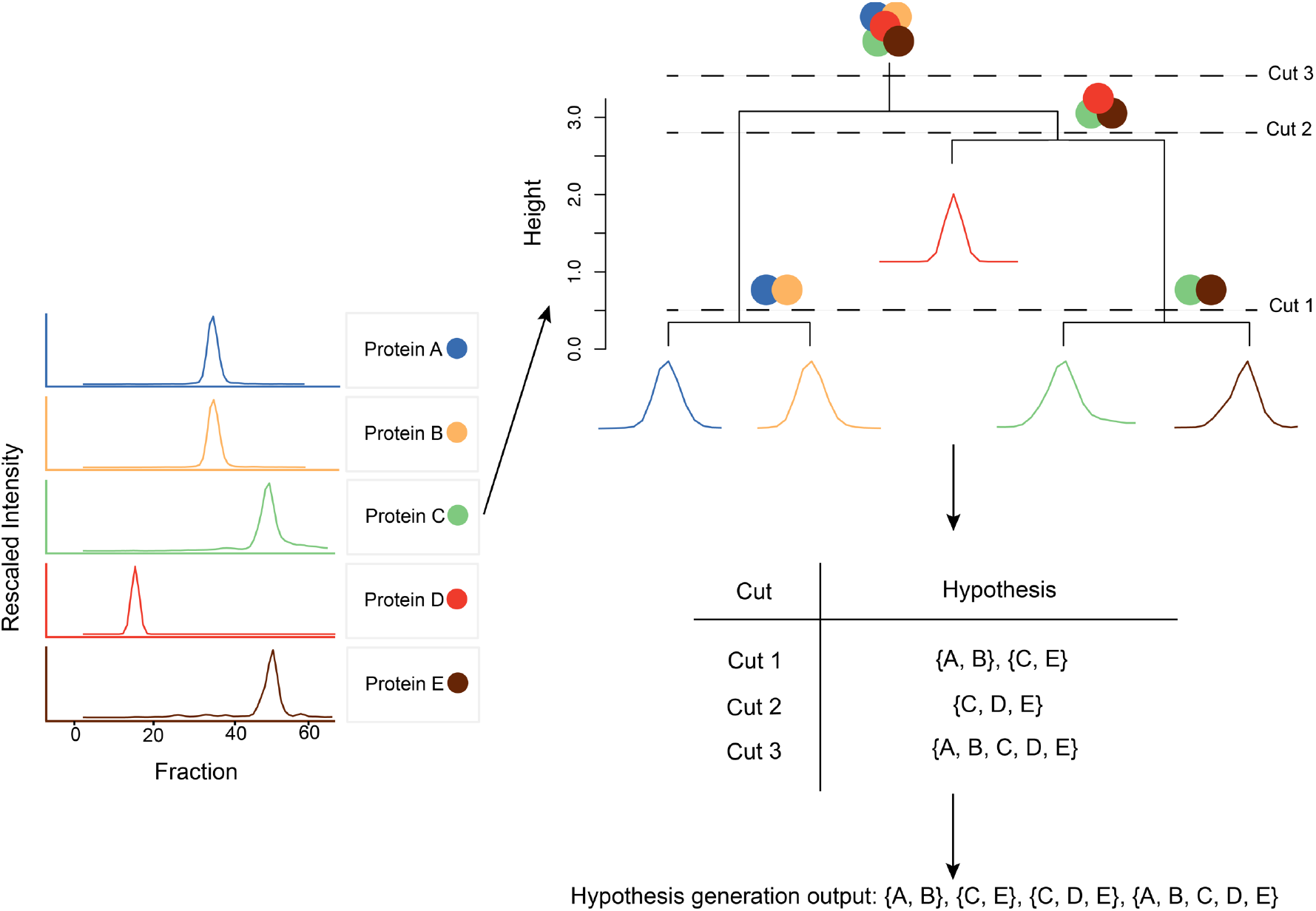
A conceptual illustration of hypothesis generation. Every protein is clustered into possible complexes using Euclidian distance clustering. Following construction of a dendrogram, all resulting clusters are retrieved by cutting at all heights (i.e. distances). This generates a comprehensive set of all possible complexes in the data.

**Supplementary Fig. S5.**
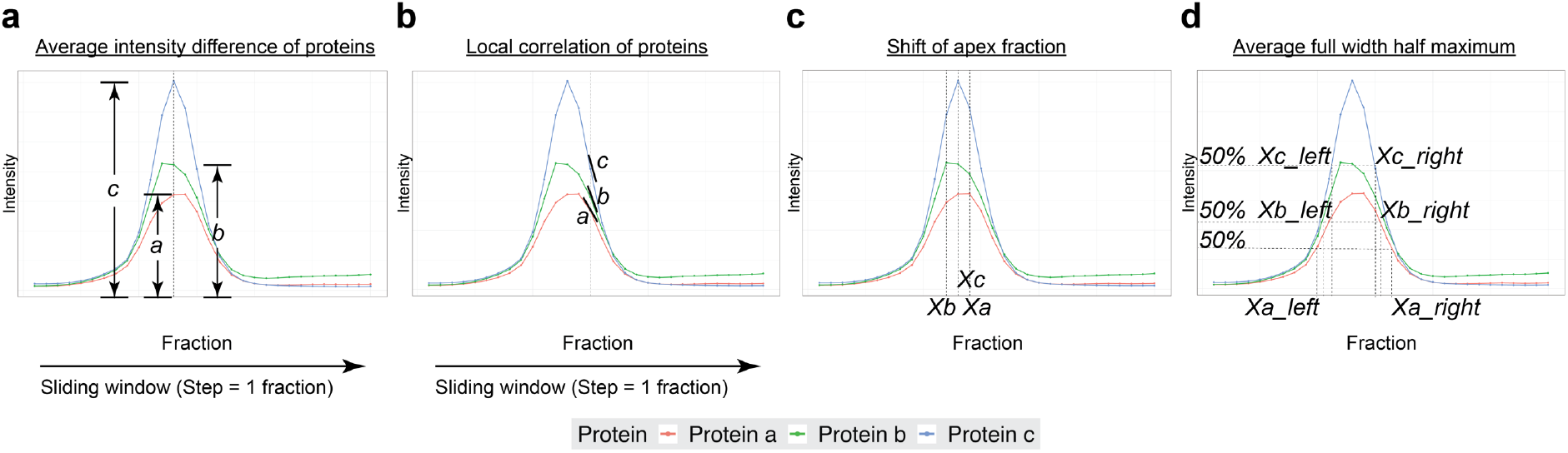
Feature calculation based on protein co-elution profiles. **a**, average intensity difference of proteins. **b**, local correlation of proteins at each fraction. **c**, shift of apex fraction. **d**, average full width half maximum.

**Supplementary Fig. S6.**
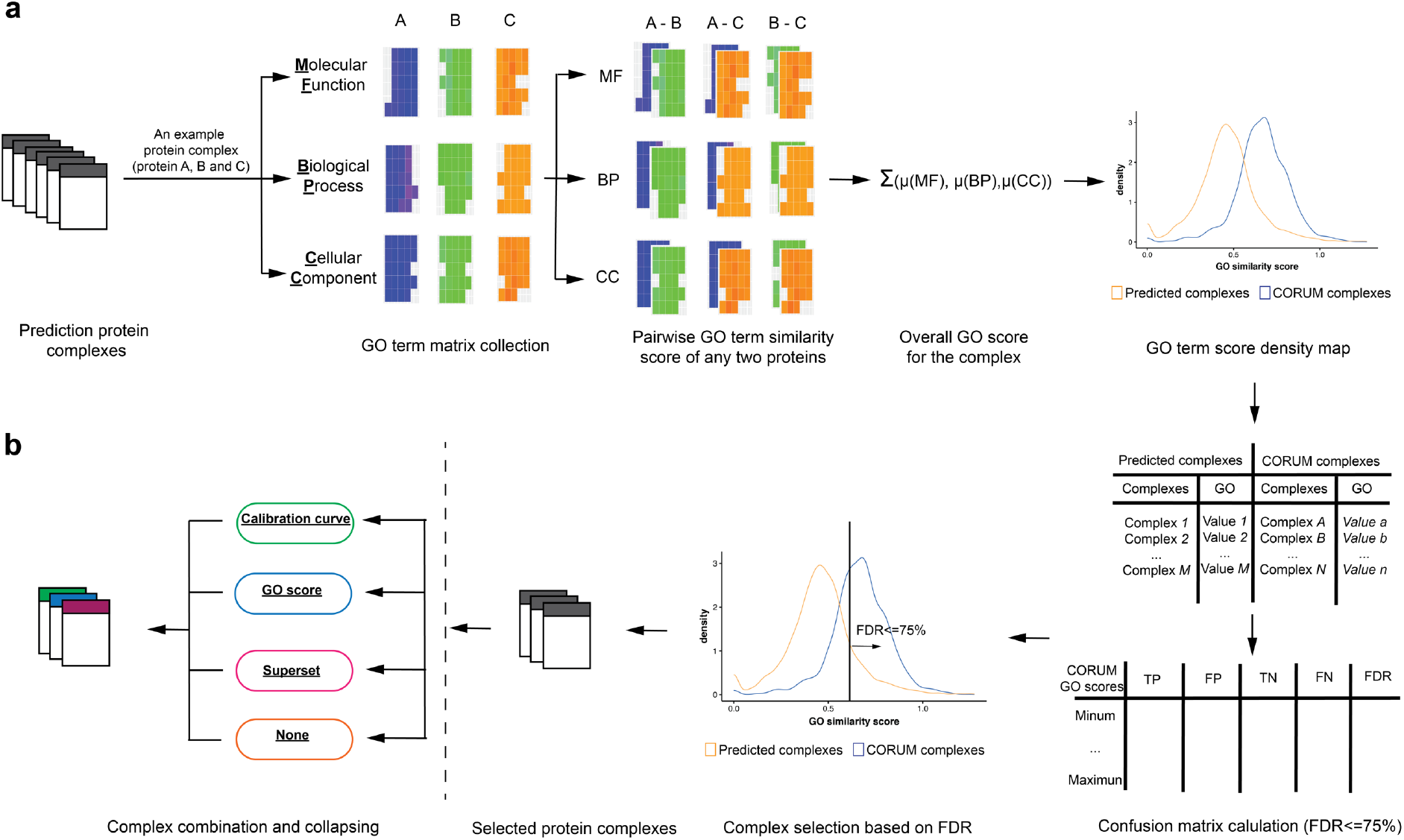
Graphical illustration of post-prediction processing. **a**, GO term score filtering. **b**, complex combination and collapsing. Positively predicted complexes either from the provided database or PCprophet are decomposed into PPI and pairwise metrics are calculated based on semantic similarity between the different ontologies which is then filtered based on a user-defined FDR threshold. Overlapping complexes are combined based on the user-defined criteria.

**Supplementary Fig. S7.**
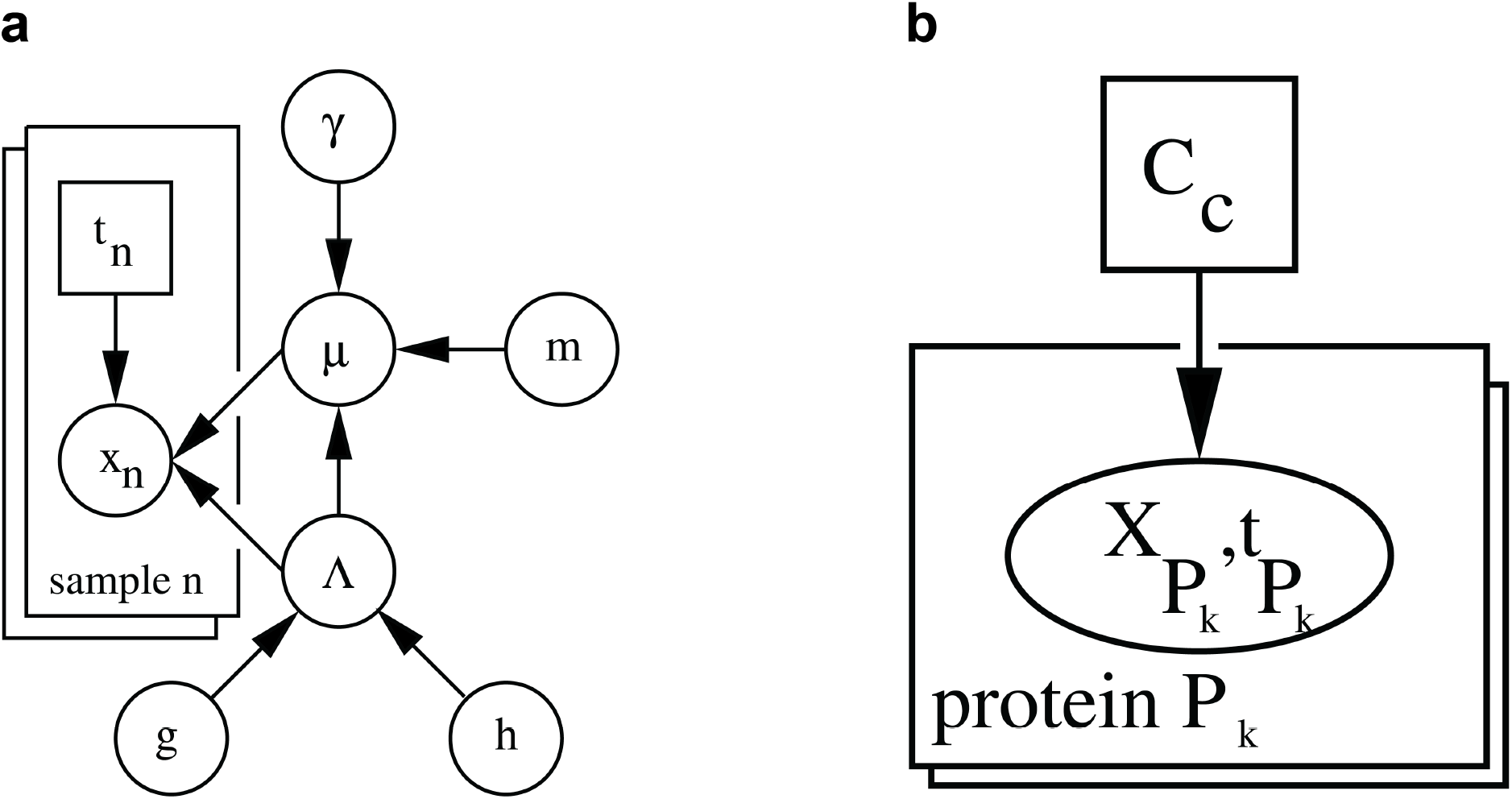
Using Bayesian inference to analyse the difference of protein complexes across conditions. **a**, An analytically tractable model for inferring differentially regulated proteins. Variable *μ* denotes the mean of a multivariate Gaussian distribution over the protein abundance vector *X_n_* which depends on the phenotype state tn. The prior over *μ* is a Gaussian distribution and parameterized by *m* (the prior location), Λ and *γ* (together specifying the precision of the Gaussian distribution). Variable *Λ* acts also as a precision matrix in the Gaussian *p*(*x_n_|μ*, Λ). The prior over Λ is a diagonal Wishart distribution (a product of Gamma distributions) which is parameterised by the hyper parameters *g* and *h*. The conditional distribution *p(μ*, Λ\g, h, m, γ) which is represented by this DAG is referred to as Normal-Wishart distribution and allows for an analytical calculation of Bayes factors. **b**, A probabilistic model for inferring differentially regulated protein complexes. Variable *Cc* denotes the state of differential regulation of a protein complex as binary variable. The elliptic node ***X**_P_k__, **t**_P_k__* represents the marginal likelihood of the protein retention profiles ***X**_P_k__* of protein *P_k_* ∈ *C_c_* in dependence of the phenotype characterization ***t**_P_k__* as they arise from Equation (15) for *C_c_* = 1 and from Equation (16) for *C_c_ =* 0. For calculating the state of differential regulation of protein complex *Cc* we make thus a conditional independence assumption among all contributing marginal likelihoods.

**Supplementary Fig. S8.**
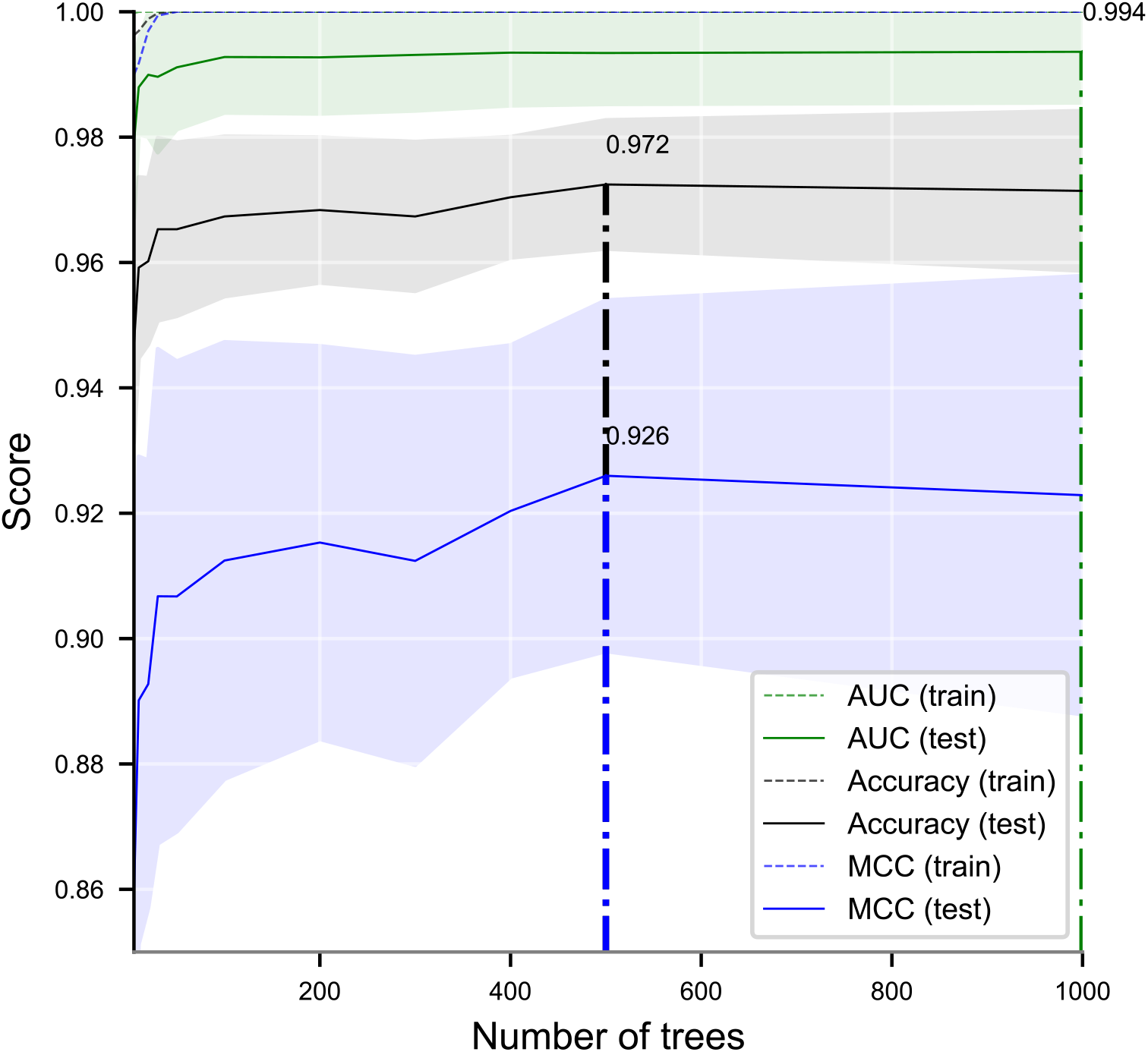
Determination of the number of trees in the RF model using three performance evaluation measures, including accuracy, AUC and MCC, via stratified 10-fold cross-validation on the entire dataset.

**Supplementary Fig. S9.**
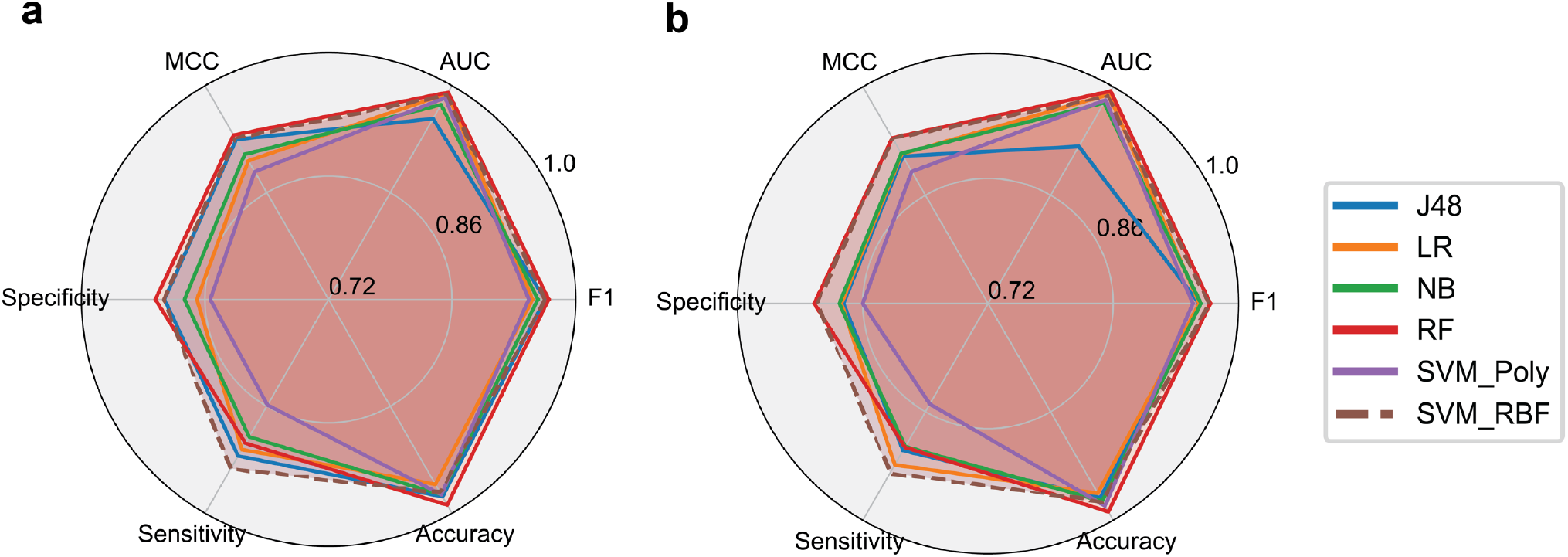
Radar plots demonstrating the prediction performance of PCprophet via five-fold cross-validation. Performance parameters are AUC, accuracy, F1, MCC, sensitivity and specificity. The analysis was based on the HEK293 dataset^10^ using **a**, equal numbers of positives and negatives, which were randomly selected 100 times and **b**, positives and all negatives. Coloured lines show the performance measures of different machine-learning algorithms namely J48 decision tree (J48), linear regression (LR), Naïve Bayes (NB), Random Forest (RF) and Support Vector Machine with either polynomial kernel (SVM_Poly) or radial basis function kernel (SVM_RBF)

## Supplementary Tables

**Supplementary Table S1.** AUC values evaluating the similarity between ground-truth networks and prediction

| AUC | Software | ΔAUC |
| --- | --- | --- |
| **PCprophet** | 0.096 | 0.013 |
| **EPIC_SVM** | 0.390 | 0.282 |
| **EPIC_RF** | 0.175 | 0.067 |
| **CCprofiler** | 0.122 | 0.014 |
| **BioPlex** | 0.040 | 0.068 |
| **STRING** | 0.137 | 0.029 |
| **CORUM** | 0.109 | 0 |

**Supplementary Table S2.** Detailed descriptions of the datasets applied for training and evaluating PCprophet

| Dataset | Cell line | Species | Data acquisition method | Separation technique |
| --- | --- | --- | --- | --- |
| HEK293<sup>10</sup> | HEK293 | *Homo sapiens* | SWATH | SEC (Size Exclusion Chromatography) |
| HeLa mitosis and interphase<sup>17</sup> | HeLa CCL2 | *Homo sapiens* |  |  |
| DDA-SILAC HeLa<sup>18</sup> | HeLa | *Homo sapiens* | DDA-SILAC |  |

**Supplementary Table S3.** An example demonstrating the complex collapsing step using three protein complexes

| Complex | Subunit | GO score | Calibration MW | Apparent MW |
| --- | --- | --- | --- | --- |
| PC1 | A, B | 0.5 | 150,000 | 100,000 |
| PC2 | A, B, C | 0.9 | 150,000 | 130,000 |
| PC3 | A, B, C, D | 0.7 | 150,000 | 160,000 |

**Supplementary Table S4.** Prediction performance of PCprophet on manually annotated HEK293 datasets via 5-fold cross-validation using different numbers of negatives

|  | AUC | Accuracy | F1 | MCC | Specificity | Sensitivity |
| --- | --- | --- | --- | --- | --- | --- |
|  | Using equal numbers of positives and negatives |  |  |  |  |  |
| **J48** | 0.957 | 96.531% | 0.929 | 0.906 | 0.978 | 0.925 |
| **LR** | 0.990 | 95.102% | 0.902 | 0.869 | 0.962 | **0.917** |
| **NB** | 0.975 | 95.714% | 0.910 | 0.883 | 0.976 | 0.900 |
| **RF** | **0.991** | **96.939%** | **0.935** | **0.916** | **0.989** | 0.908 |
| **SVM_POLY** | 0.984 | 94.694% | 0.887 | 0.854 | 0.976 | 0.858 |
| **SVM_RBF** | 0.988 | 96.531% | 0.931 | 0.907 | 0.973 | 0.942 |
|  | Using positives and all negatives |  |  |  |  |  |
| **J48** | 0.923 | 95.603% | 0.910 | 0.882 | 0.971 | 0.910 |
| **LR** | 0.990 | 95.622% | 0.913 | 0.884 | 0.965 | 0.929 |
| **NB** | 0.979 | 95.796% | 0.914 | 0.886 | 0.975 | 0.905 |
| **RF** | **0.994** | **97.268%** | **0.934** | **0.926** | **0.989** | 0.906 |
| **SVM_POLY** | 0.983 | 94.896% | 0.891 | 0.860 | 0.981 | 0.850 |
| **SVM_RBF** | 0.988 | 96.684% | 0.933 | 0.911 | 0.976 | **0.939** |

## Notes

### Competing Interest Statement

The authors have declared no competing interest.

